# The ΔF508 CFTR defect: molecular mechanism of suppressor mutation V510D and the contribution of transmembrane helix unraveling

**DOI:** 10.1101/2020.04.19.049338

**Authors:** Kailene S. Simon, Karthi Nagarajan, Ingrid Mechin, Caroline Duffy, Partha Manavalan, Steve Altmann, Aliza Majewski, Joseph Foley, J. Stefan Kaczmarek, Scott Bercury, Matthew Maderia, Brendan Hilbert, Joseph D. Batchelor, Robin Ziegler, Jeffrey Bajko, Michael Kothe, Ronald K. Scheule, Anil Nair, Gregory D. Hurlbut

## Abstract

Cystic fibrosis (CF) results from mutations within the gene encoding the Cystic Fibrosis Transmembrane Conductance Regulator (CFTR), a transmembrane chloride channel found on the apical surface of epithelial cells. The most common CF-causing mutation results in a deletion of phenylalanine 508 (ΔF508-CFTR), a residue normally found within the NBD1 domain. Loss of F508 causes NBD1 to be less thermodynamically stable and prevents proper tertiary folding of CFTR. As a result, CFTR is not properly trafficked to the cell surface. Recently, progress has been made towards the development of small molecule “correctors” that can restore CFTR tertiary structure and stabilize the channel to overcome the instability inherent in ΔF508-CFTR. However, the resultant improvement in channel activity has been modest, and the need for potent correctors remains. To fully inform such efforts, a better understanding of the molecular pathology associated with ΔF508-CFTR is required. Here we present a comprehensive study of the impact of F508 deletion on both purified NBD1 and full-length CFTR. Through the use of homology modeling, molecular dynamics simulations, mutational analysis, biochemical, biophysical and functional characterization studies, we obtained insight into how the ΔF508 mutation may lead to helical unraveling of transmembrane domains 10 and 11 (TM10, TM11), and how the known suppressor mutations V510D and R1070W, as well as novel second site suppressor mutations (SSSMs) identified in this work, may act to rescue ΔF508-CFTR maturation and trafficking.

## Introduction

Cystic Fibrosis (CF) is a genetic disease caused by mutations to the gene that encodes cystic fibrosis transmembrane conductance regulator (CFTR), a chloride channel found in the apical membrane of epithelial cells. Its basic topology consists of two transmembrane domains (TMD1 and TMD2), each with six transmembrane (TM) α-helices, two cytoplasmic nucleotide-binding domains (NBD1 and NBD2) and a unique regulatory (R) domain with multiple phosphorylation sites (1–3). CFTR gating is regulated by PKA-dependent phosphorylation of the R-domain and ATP-binding and dimerization of the cytoplasmic NBDs, resulting in an outward-facing, open conformation of the channel that facilitates passive Cl^-^ transport down its concentration gradient (4).

Roughly 70% of CF cases worldwide result from a deletion of F508 (ΔF508-CFTR) within NBD1. Loss of this aromatic residue reduces NBD1 stability, as demonstrated by a decreased (6–8°C) unfolding transition temperature (Tm) of purified NBD1 (5). In the context of full-length CFTR, the loss of F508 and this resulting NBD1 instability, disrupts the interface between NBD1 and intracellular loop 4 (ICL4) of the second transmembrane domain (TMD2) (6). Together, these defects diminish proper CFTR folding, apical trafficking, and channel function (6). ΔF508-CFTR is instead largely retained in the endoplasmic reticulum and targeted for subsequent degradation (1,7). While the impact of ΔF508 on CFTR folding and stability has been known for some time (7,8), the mechanisms by which F508 loss destabilizes the channel have only more recently been described (1,7,9).

Previously reported crystal structures and hydrogen-deuterium exchange (HDX) analysis of both wild type-(WT) and ΔF508-NBD1 have provided evidence that F508 loss does not impact NBD1 structure globally, but instead results in localized solvent exposure of the V510 loop (1,10,11). This result is consistent with *in silico* NBD1 models, which suggest the same localized impact at the V510 loop (12–14). Because a full-length ΔF508-CFTR structure has not yet been published, much of what is known about the compromised molecular interactions and structural defects at the NBD1:ICL4 interface is the product of molecular modeling. Such work has provided valuable insight into the local environment of the V510 loop, as well as its altered positioning with respect to ICL4 as a consequence of the mutation (15,16). CFTR homology modeling also has predicted that the loss of the large, aromatic ring structure of F508 leaves a hydrophobic cavity at the inter-domain interface of NBD1 and ICL4 (17).

A series of CFTR mutations have been identified, in part through patient genotyping, which significantly reduce the physiologic impact of ΔF508 when these are simultaneously present. Such CFTR second site suppressor mutations (SSSMs) result in improved CFTR trafficking and function, and a milder disease phenotype (18,19). Among SSSMs, I539T, G550E, R553Q, and R555K are located within NBD1. These mutations have been shown to increase the thermal stability of the domain, but do not (in models) appear to reestablish the NBD1: ICL4 interface that is lost due to ΔF508 (3). However, when ΔF508 is coupled with SSSMs that act as interface correctors between NBD1 and ICL4, a vast improvement can be seen in ΔF508 CFTR global assembly and channel function (5,18,19). One such example, R1070W, may drive ΔF508 phenotype suppression by replacing the missing aromatic ring of F508 with a tryptophan at the ICL4 interface (17,20). Another example, which was first identified during an attempt to create Cys-less CFTR (21) and supported with homology modeling and biochemical characterization shortly thereafter, is V510D (15,20), which is thought to restore the NBD1:ICL4 interface by creating a salt bridge with R1070 on ICL4. The half-life of ΔF508/V510D-CFTR at the cell surface is reported to be similar to that of wild type CFTR, 5-10 fold longer than that of ΔF508 (20). In addition to correcting the NBD1:ICL4 interface, other possible interactions may be occurring within this ΔF508/V510D CFTR double mutant that may be critical to our understanding of how such mutations act to suppress the ΔF508 phenotype.

While it is well established that suppressor mutations can rescue ΔF508-CFTR trafficking, the mechanism driving this rescue is only partially understood. This work was therefore undertaken to accomplish three major goals: i) to construct and validate a molecular model of ΔF508-CFTR in the absence of a full-length structure, ii) to use this model to gain insight into the effects of the ΔF508 mutation on CFTR conformational dynamics, and iii) to determine the mechanism(s) by which second-site suppressor mutations may be influencing ΔF508-CFTR conformational dynamics.

To accomplish these goals, we generated homology models of full-length WT and ΔF508-CFTR +/− the SSSMs V510D and R1070W and performed 1 microsecond molecular dynamics (MD) simulations for each. These simulations provided key insights into how the ΔF508 defect may be impacting CFTR structural integrity for both NBD1 and full-length CFTR (without the regulatory domain) and led to several testable hypotheses regarding ΔF508 destabilization of full-length CFTR. They also identified residues - most notably K564 - that may play a critical role in the mechanism by which V510D both stabilizes NBD1 and improves ΔF508-CFTR pathology. By then creating full-length WT and ΔF508-CFTR constructs containing certain SSSMs or other point mutations, we gained additional new insights into how the V510D suppressor mutation has a corrective effect on ΔF508. Moreover, we determined the structure of ΔF508/V510D-NBD1 by X-ray crystallography. By comparing the crystal structures of ΔF508-NBD1 and ΔF508/V510D-NBD1, we determined what structural changes and interactions resulted within NBD1, either at the V510 loop or other regions of ΔF508-NBD1, that drive V510D stabilization of this soluble domain. In an effort to improve the stability of the TMD2 domain, we introduced charged residue substitutions along TM10 and TM11 with a view to create potentially stabilizing salt bridges within TMD2. We then evaluated the impact of these mutations on CFTR maturation and trafficking, both alone and in the context of additional mutations within NBD1 and ICL4. The experimental data (expression, trafficking and functional) obtained from these mutants support the hypothesis that a contributing factor of the ΔF508 defect may be the helical instability of the second transmembrane domain.

## Results

### CFTR Homology Model and Molecular Dynamics Simulation

Using the homologous ABC transporter Sav1866 (22) (PDB ID: 2HYD) as a model, we generated novel molecular models of WT and ΔF508-CFTR (Figure S1) (22). Although multiple CFTR structures have been published recently (23–27), this was the best template available when we started our computational and experimental studies. For the WT CFTR model, we replaced the NBDs of Sav1866 with CFTR NBD1 and NBD2 crystal structures (PDB IDs: 2PZE and 3GD7, respectively). To generate the ΔF508-NBD1 model we inserted the ΔF508-NBD1 crystal structure (PDB ID: 2PZF) (1,10). Relative positioning of the NBDs was based on Sav1866 and NBD1 homodimer structures (1,22).

To evaluate the accuracy of our WT CFTR model, we compared its structure to that of the recently published outward-facing, open CFTR conformation obtained by cryo-EM, paying particular attention to NBD1, NBD2 and the intracellular loops (ICLs), as those domains are the focus of this study. Figure 1 shows that structural alignment of those regions of our model with those of the experimentally-derived cryo-EM data (PDB ID: 6MSM, (27)) show high structural similarity with an overall root-mean-square deviation (RMSD) for C-alpha atoms of 3.76Å. Both NBDs display a high level of correlation between the model and cryo-EM structure. Alignment of NBD1 from our WT-CFTR model (based on NBD1 crystal structure PDB ID: 2PZE) and the cryo-EM structure returned an RMSD of 1.14 Å. Alignment of NBD2 from our model (based on the CFTR-NBD2-maltose binding protein fusion complex crystal structure; PDB ID: 3GD7) with the analogous cryo-EM domain gave an RMSD of 1.916 Å.

**Figure 1.**
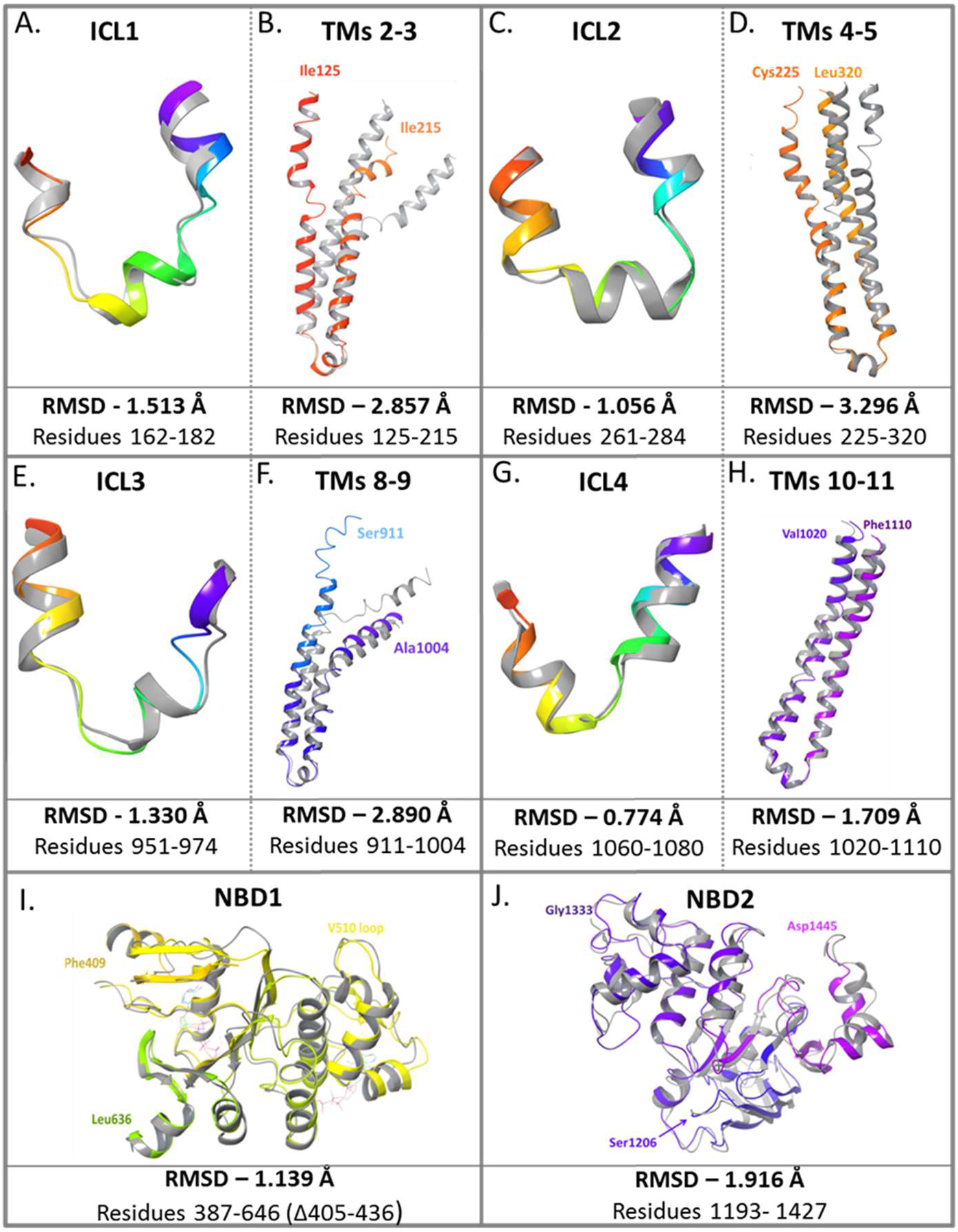
Structural comparison and corresponding RMSD values of ICL and NBD regions for the human WT-CFTR homology model (colored ribbon structures) and the published cryo-EM structure of CFTR (grey ribbon structures, PDB ID: 6MSM) (27). Both structures represent CFTR in an outward-facing conformation. Structural similarities between the two models are highlighted for each intracellular loop (ICL) (panels A, C, E and G) displayed alongside the corresponding pair of transmembrane helices that comprise that ICL (B, D, F, and H, respectively), as well as an overall comparison of NBD1 (panel I) and NBD2 (panel J).

Figure 1 also provides RMSD values comparing ICL and TM domains of our full-length WT-CFTR homology model with those of the corresponding cryo-EM CFTR structure. Positioning in relation to the TMs would likely be slightly different, given that our model is based on Sav1866. Panels B, D and F highlight greater differences in TMs 3, 5 and 8, respectively, between the homology model and cryo-EM structure. In our homology model, the helices appear to be slightly straighter, while the cryo-EM model displays a helix-loop-helix structure for TMs 3 and 8. In all cases, the alignment appears different for one or both TMs that make up the pair. This may be due to the discrepancies between the channel pore size of Sav1866, which is larger to facilitate the passage of bigger molecules, and CFTR, which need only allow for the passage of ions. Importantly, the ICL4-flanking TMs 10 and 11 which are a focus of our study show good alignment between our model and published CFTR structures.

Additional ΔF508-CFTR variant models were generated by incorporating the known SSSMs V510D and R1070W. Using these models, we built corresponding ~160,000 atom explicit membrane/explicit solvent systems with which to conduct 1 μsec MD simulations (Figure S2).

#### Quality Analysis for Molecular Dynamics Simulations

A quality analysis comparison was completed for the MD simulations (Figure S3). Macroscopic properties (potential and kinetic energy, pressure, volume, temperature) of the models evaluated over time confirmed that stable simulations were performed. All physical properties evaluated showed an initial relaxation of the models followed by stabilization.

In analyzing these MD simulation outcomes, we looked for specific and testable changes in secondary structure (Figures 2–4), including solvent exposure, residue geometry, and the existence and conservation of salt bridges between selected residues in each of the models (Figure S4). Figure 2 shows that significant conformational changes were seen within TMD2 for ΔF508 and ΔF508+R1070W, which are highlighted by the reduction in structured regions from residues 1044 to 1095. When V510D is added to ΔF508-CFTR, helical structure appears to be better maintained in this region, as evidenced by a secondary structure profile more closely resembling WT. Still shots of the MD movies illustrate this helical unraveling along TMs 10 and 11, and at ICL4 that connects them (Figure 3). A more detailed view of the unraveling in the TM10 alpha helical region (residues 1028 – 1055) is shown in Figure 4. Superimposing the structures of WT at the beginning and end of the MD simulation shows little difference in conformation, while in ΔF508-CFTR, residues 1034 – 1048 become unraveled and lose their initial alpha helical conformation.

**Figure 2.**
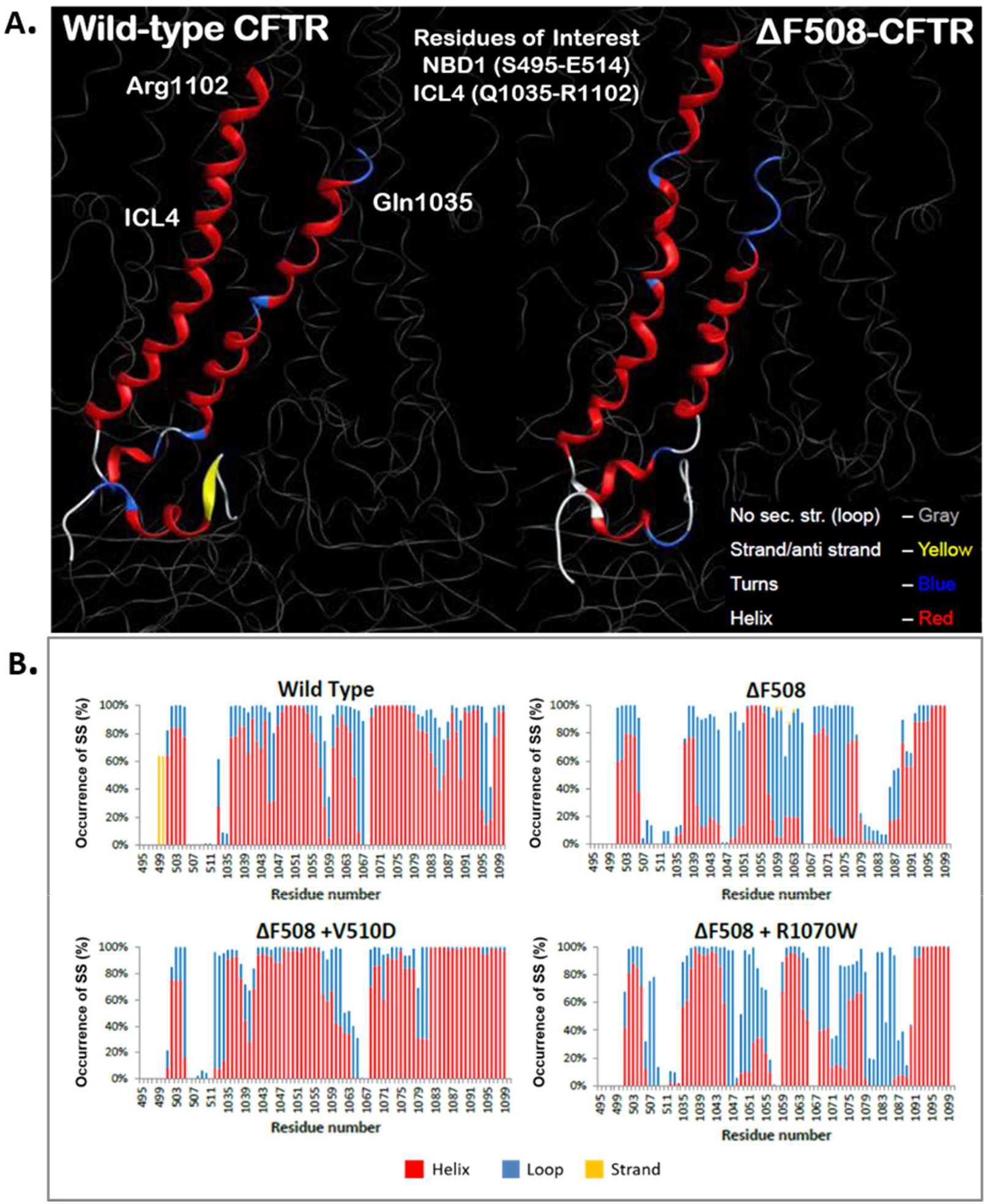
CFTR residues of interest in MD simulations. A. Regions in NBD1 and ICL4 and their corresponding residue numbers that were highlighted for differences in secondary structure throughout the course of the 1 μsec MD simulations. In the MD simulations, the models were color-coded to signify secondary structures: helix = red; strand/anti-strand = yellow; turns = blue; no secondary structure = gray. B. Occurrence percentage of secondary structural changes in models for ΔF508, ΔF508+V510D and ΔF508 + R1070W was consistent with reported stability data for these variants. A comparison of secondary structure classifications by percent occurrence for each residue of interest within the four CFTR homology models shows helical regions in red, loops in blue and strands in yellow.

**Figure 3.**
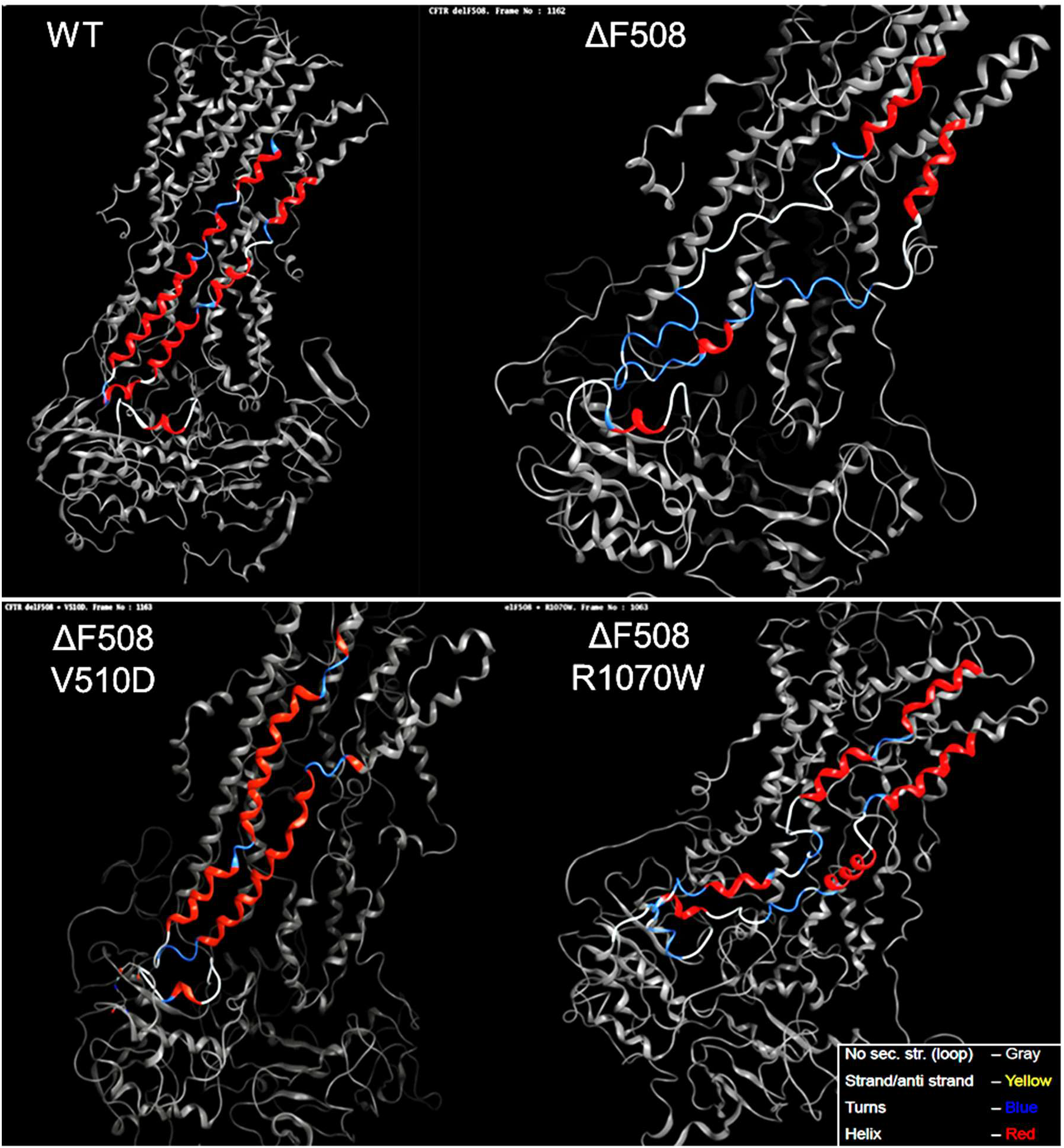
Molecular dynamics simulations of WT, ΔF508, ΔF508/V510D and ΔF508/R1070W. Still shots were taken of each of the four movies at the same point of progression in the simulation, highlighting the vast differences in helical stability across the four CFTR molecules.

Importantly, a comparison of the four (WT, ΔF508, ΔF508+V510D and ΔF508+R1070W) 1 μs MD simulations also demonstrated a strong correlation between *in silico* protein stability and published experimental data such as secondary structure stability (WT ~ ΔF508+V510D >> ΔF508+R1070W > ΔF508), and supported a proposed salt-bridge interaction between V510D and R1070 at the NBD1:ICL4 interface (28,29). These results also support previously made predictions regarding increased solvent exposure of the V510 loop in the V510D mutant (15).

### V510D Salt Bridge Interactions in ΔF508-CFTR

The MD analysis also offered several new insights concerning key residues within NBD1 that may play a role in V510D stabilization of ΔF508-NBD1. Potential salt bridges were retrieved for each charged residue from individual frames of the simulation. Residues with a high occurrence of interactions were analyzed further for their total number of salt bridge interactions over time. Frequency analysis of intra- and inter-domain salt bridge possibilities for the MD run of the ΔF508/V510D model indicated that 510D may interact with K564 and, to a lesser extent, with R487 in NBD1 (Figure S4). This previously undescribed interaction with K564 is in addition to its anticipated interaction with R1070 of the ICL4 loop.

#### K564 is Key to V510D Suppression of the ΔF508 Trafficking Defect

To evaluate experimentally the *in silico* model prediction that D510 might interact with K564 and/or R487, these residues were replaced with alanine or serine in WT, ΔF508–CFTR, and ΔF508/V510D-CFTR expression constructs to test the impact of removing a positive charge in close proximity to D510, effectively eliminating a potential stabilizing interaction for the solvent-exposed V510 loop of ΔF508-NBD1. A ΔF508/V510K/K564D construct was also created to evaluate whether ΔF508-CFTR rescue can occur if residue sidechains at 510 and 564 were swapped, potentially recapitulating the proposed K5 64/V510D salt bridge within NBD1, but without enabling the V510D:R1070 interaction to stabilize the NBD1:ICL4 interface. We assessed the impact of each mutation on ΔF508-CFTR maturation and trafficking by transiently transfecting each construct into CF patient-derived submucosal gland epithelial cells (CFSME_0_^-^) and comparing the resulting expression levels of mature, fully glycosylated CFTR (“C-band”) to those of immature, core-glycosylated CFTR (“B-band”) 48 h post-transfection and expressing results as a ratio of C/B band intensities. Since this ratio does not reflect overall changes in CFTR expression, we also generated a second set of constructs with the same point mutations that contained an in-frame fusion of horseradish peroxidase (HRP) within CFTR’s 4^th^ extracellular loop, allowing us to more quantitatively detect membrane-bound CFTR using an HRP-mediated signal. To ensure that samples in both the western blot and HRP trafficking assays were normalized for CFTR transfection efficiency, constructs were designed with a coexpressing soluble eGFP marker using the 2A bicistronic expression system (30,31). This system ensures a 1:1 ratio of CFTR and GFP protein production. Results are shown in Figure 5.

**Figure 4.**
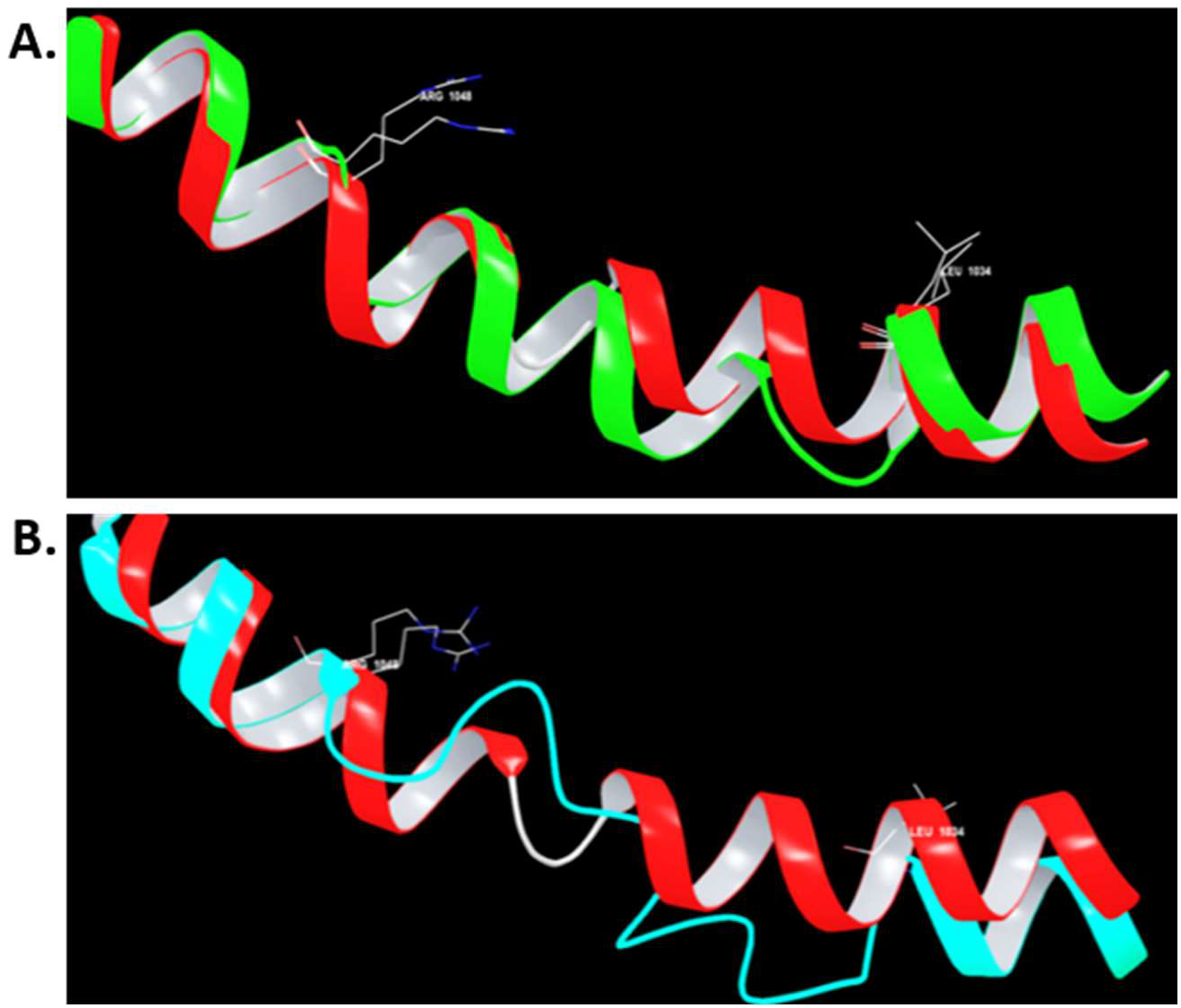
Superposition of TM10 alpha helical region (residues 1028-1055) taken from the first (red) and last frame (green/teal) of MD simulations for WT (A) and ΔF508 (B) models of full-length CFTR. The simulation time between first and last frames is 1 μsec. (A) MD simulation for WT-CFTR suggests that the alpha-helical conformation of TM10 does not change over the course of the simulation; however, a slight adjustment in position may occur. RMS deviation between these two conformations is 1.55 Å. (B) A comparison of conformations for ΔF508-CFTR TM10 region between the first frame (red helix) and last (cyan) highlights changes in conformation between residues 1034 and 1048, which is unwound and loses α-helical conformation. RMS deviation between these two conformations is 2.38 Å.

**Figure 5.**
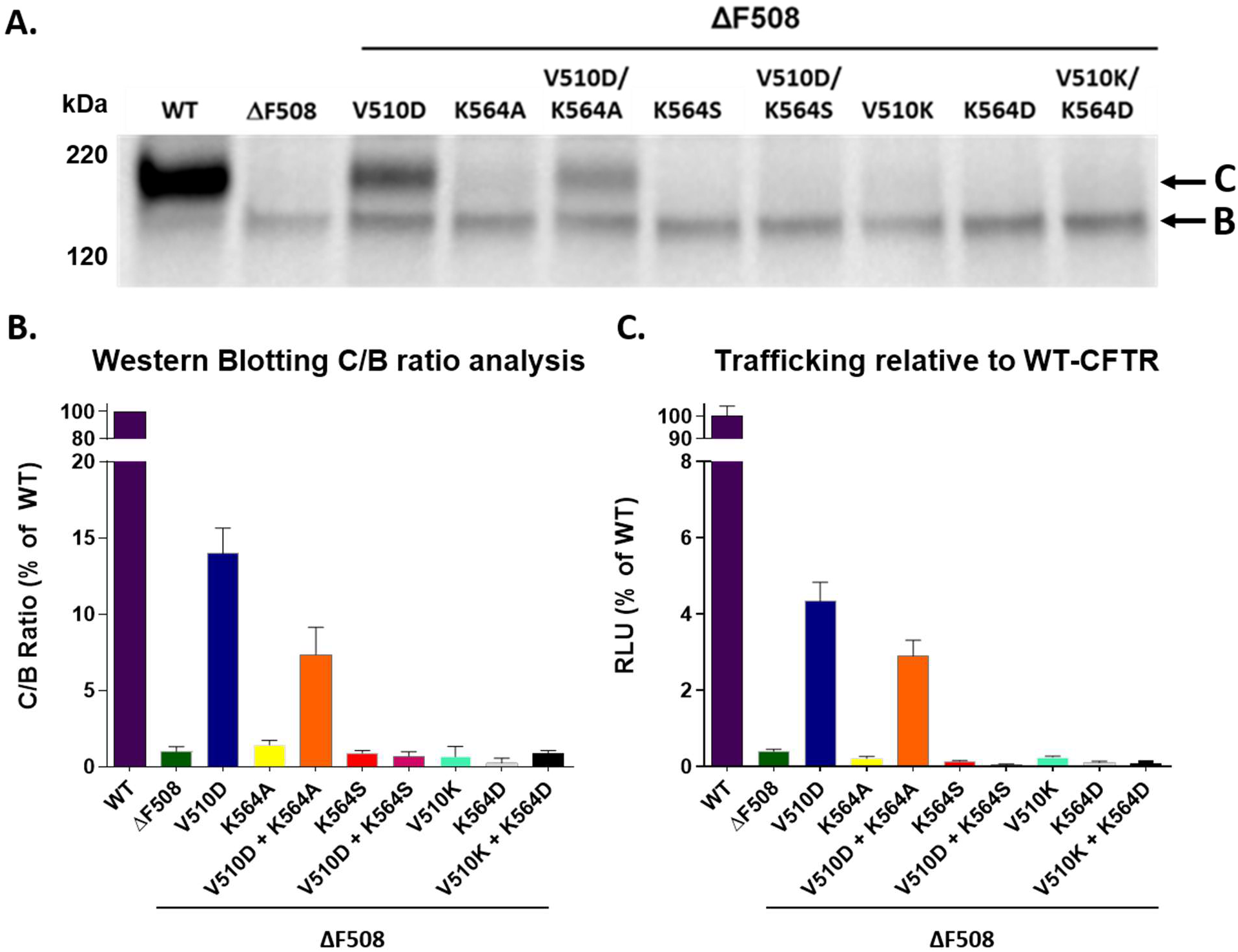
The role of K564 on V510D-mediated suppression on the ΔF508 trafficking defect. Immunoblotting analysis (A) and C/B ratio values (B) comparing WT and ΔF508-CFTR to V510D and K564 mutants, showing partial CFTR rescue (as indicated by the increase in “C-band”) when V510D is added to ΔF508. This rescue effect is diminished when the K564A mutation is added to V510D. K564S and residue-swap mutations containing K564D and V510K completely eliminate CFTR trafficking. (C) An HRP reporter assay was used to measure trafficking of the CFTR variants to the cell surface. Data are displayed as RLU values relative to WT-CFTR.

Initial analysis was performed on WT-CFTR constructs containing the K564A mutation +/− V510D prior to evaluation on ΔF508-CFTR to confirm that the K564 mutation had no deleterious effect on CFTR maturation (Figure S5). Figure 5 shows that immunoblotting for CFTR displayed no reduction in levels of fully glycosylated CFTR when mutations were added, and an increase in overall amounts of mature CFTR was detected. While some rescue of ΔF508-CFTR is evident when V510D is present, (restoring trafficking to 5-10% of WT levels), our data suggest that removing charged residue K564 disrupts V510D rescue of the ΔF508 trafficking defect. Indeed, when K564 was mutated to an alanine on a ΔF508/V510D background, only half of the rescue effect was preserved, suggesting that the impact of V510D rescue within NBD1 may rely on the presence of this lysine. Interestingly, when K564 was replaced with a serine, or when D510 and K564 were transposed, trafficking in both the HRP and western blot assays decreased to ΔF508 levels.

### V510D Impact on the Structure and Stability of NBD1

Given the inherent instability of NBD1 in its native form, much has been done to identify both point mutations and peptide deletions that improve yield and solubility of the purified protein (1,6,32,33). As described above, such NBD1 mutations include known CFTR SSSMs like G550E, R553Q, and R555K. NBD1-stabilizing modifications also include removing the NBD1 37-residue regulatory insertion (RI, residues 402-438) with or without a further truncation of the majority of the 38-residue regulatory extension (RE, residues 638-676), either of which results in improved NBD1 thermostability as well as increased CFTR trafficking and half-life in the presence or absence ofΔF508. (6). To that end, in an effort to understand better how the V510D suppressor mutation impacts folding and stability of the NBD1 subdomain independent of ICL4, a series of assays was performed on purified ΔRI- and ΔRI/ΔRE-NBD1 that included K564A/S mutations +/− V510D, and compared to the ΔF508 and WT versions.

#### Replacing K564 Decreases NBD1 Stability in the Presence and Absence of V510D

To evaluate each NBD1 variant for changes in thermostability, two complimentary, label-free thermal shift assays were used. The first, differential static light scattering (DSLS), measures heat-induced changes in light scattering to determine protein aggregation temperature (T_agg_). The second, nano differential scanning fluorimetry (nanoDSF), records changes to the intrinsic tryptophan (and to a lesser extent, tyrosine) fluorescence profile with increasing temperatures as a measure of thermal protein denaturation. The temperature at which the denaturation transition occurs is defined as the inflection point of the fluorescence shift (Tm).

Results from both assays are consistent and show that the V510D suppressor mutation increases the ΔF508-NBD1 T_m_ by 2-3.5°C (Figure 6). Mirroring results in the immunoblotting and HRP-trafficking assays for fulllength CFTR variants, the K564A mutation minimizes the stabilizing effect of V510D on the ΔF508 background, reducing its T_m_ by about 2°C, which represents a roughly 60% loss in V510D stabilization. Similarly, the K564S mutation eliminates the V510D rescue effect, decreasing both the melting and aggregation temperatures to below that of ΔF508-NBD1. When compared to ΔF508 alone, K564S/V510D/ΔF508-NBD1 has a ΔT_m_ of −1.37 ± 0.01°C (ΔRI/ΔRE) and −1.68 ± 0.025°C (ΔRI), and a ΔT_agg_ of −1.6 ± 0.20°C (ΔRI/ΔRE) and −0.31± 0.16°C (ΔRI).

**Figure 6.**
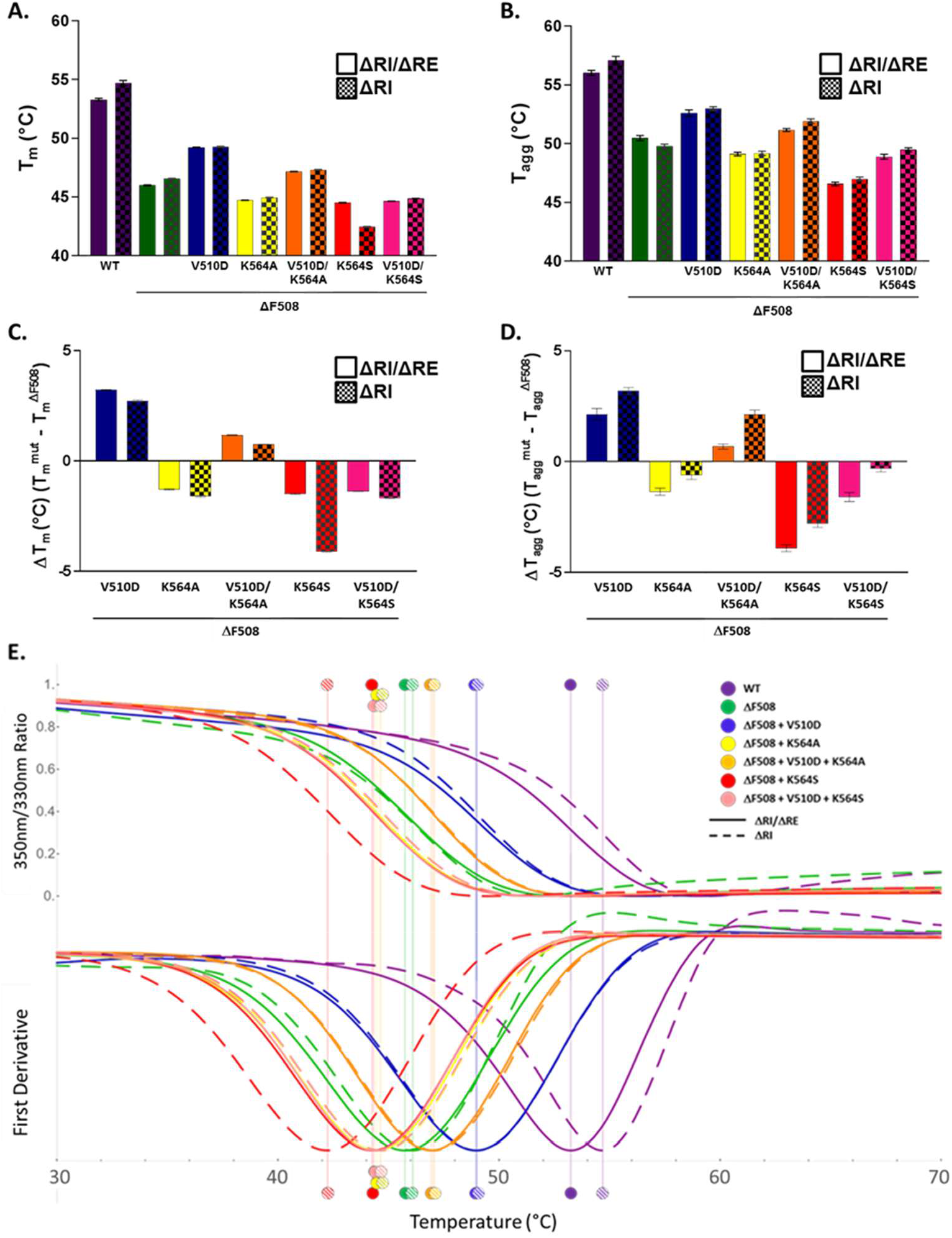
Removing K564 decreases NBD1 thermal stability in the presence and absence of V510D as indicated by DSLS and nanoDSF thermal shift assays. Purified NBD1 protein (ΔRI or ΔRI/ΔRE) was evaluated for changes in aggregation temperature (T_agg_) and melting temperature (T_m_). When point mutations K564A (yellow) and K564S (red) were made to ΔF508-NBD1 (green), a reduction of both T_agg_ and T_m_ was seen, which was partially restored by adding V510D in all cases (as indicated by blue, orange and pink). The K564S mutation resulted in a net loss of both T_agg_ and T_m_ as compared to ΔF508-NBD1, regardless of whether V510D was added. (Panels A/B) Overall T_m_ and T_agg_ (displayed as °C) for NBD1 mutants; (C/D) changes in thermal stability (ΔT_agg_ and ΔT_m_) as compared to ΔF508-NBD1; (E) Melting temperature curves for NBD1 mutants obtained by nano Differential Scanning Fluorimetry (nanoDSF). Curves are displayed as a ratio of fluorescent signals measured at 350 nm and 330 nm (top) and the first derivative of each (bottom), resulting in an inflection point indicating the T_m_ of each protein

Consistent with previously published data (6), the addition of the ΔRE truncation did not appear to contribute to NBD1 thermostability when compared to ΔRI removal alone in this set of assays, with the exception of the ΔRI/ΔF508/K564S-NBD1 protein in the nanoDSF assay, which exhibited a Tm of 2.1°C lower than its ΔRI/ΔRE counterpart. Additionally, reported T_agg_ values for all mutations were an average of 3.6°C or 3.9°C higher for ΔRI/ΔRE or ΔRI samples, respectively, when compared to their T_m_ values, likely owing this difference to the distinct phase change between native, denatured and aggregated monomeric NBD1 protein previously reported by Protasevich *et al.* (6).

#### The ΔF508 NBD1/V510D Crystal Structure Confirms the D510 – K564 Salt Bridge

While full-length CFTR trafficking and NBD1 stability data provided support for the key role of K564 in V510D-mediated rescue of ΔF508-CFTR, identifying the presence of the predicted salt bridge between K564 and V510D required a structural approach. To that end, we used crystallography to analyze changes in ΔF508-NBD1 when V510D is present. (See Supplemental Table 2 for ΔF508/V510D-NBD1 structure determination/refinement statistics.)

As has been shown previously (1), deleting F508 and the resultant shortening of the adjoining loop results in a more solvent-exposed position for V510 (Figure 7). Introducing the V510D mutation in the ΔF508 context results in a side chain rotation of D510 towards the main body of NBD1 and formation of a salt bridge with K564, whose side chain adopts an alternative rotamer from that observed in other CFTR NBD 1 structures, where the amino moiety is hydrogen-bonded to the backbone carbonyl of I488. However, when V510D is present, K564 positions the residue side chain facing toward the aspartic acid, creating a salt bridge. There are two molecules in the asymmetric unit in our crystal form. In one of them, we observe partial occupancy for both the original rotamer of K564, and the one that allows for K564-V510D salt bridge formation, while in the second molecule K564 exclusively adopts the new rotamer that allows for interaction with D510 (Figure 7). No interaction appears to exist between R487 and V510D; instead R487 is involved in a crystal contact (data not shown).

**Figure 7.**
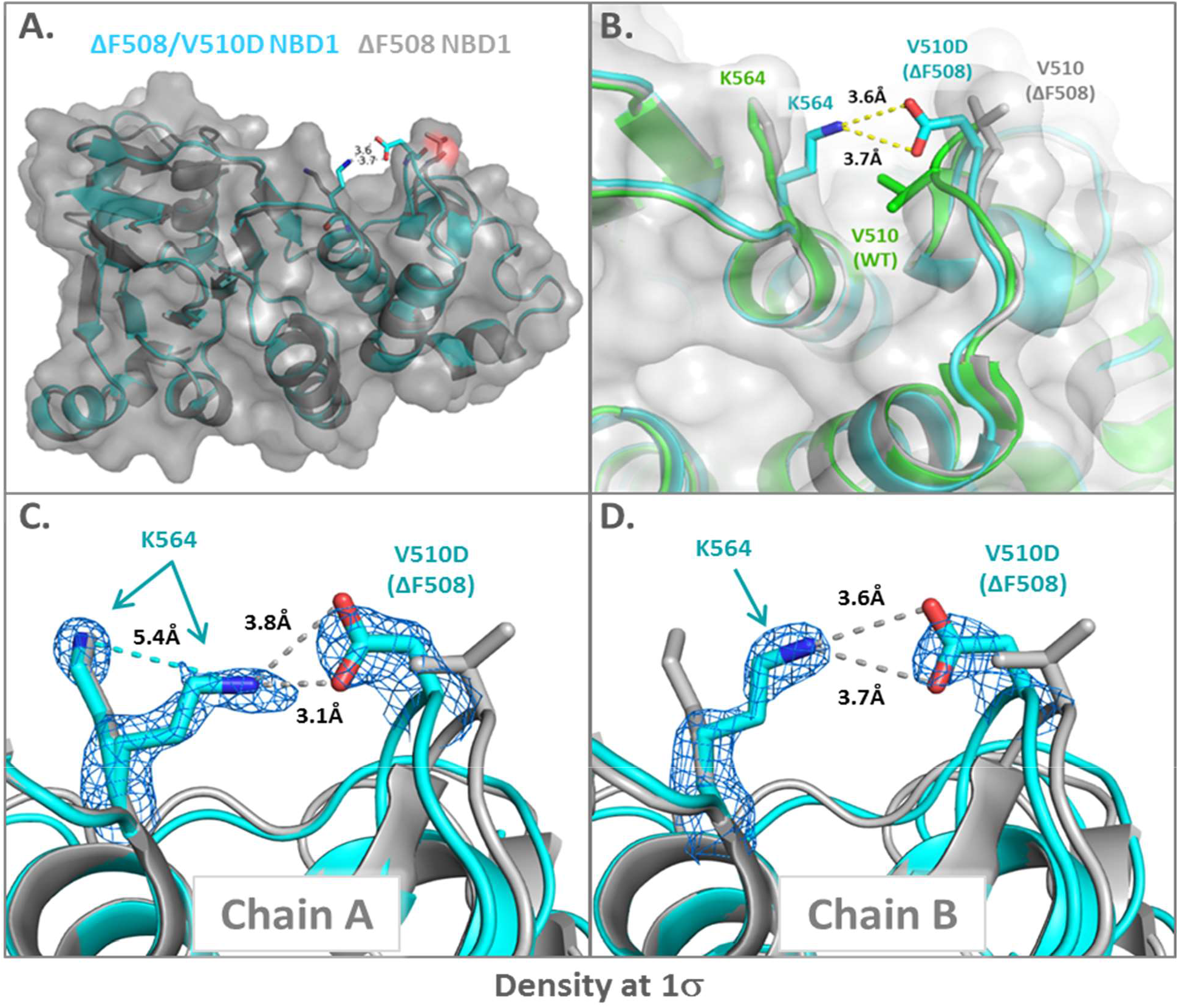
The ΔF508/V510D NBD1 crystal structure confirms the D510-K564 salt bridge. Panel A. NBD1 ΔF508/V510DNBD1 structure solved to 1.86Å resolution (shown in blue) is aligned with ΔF508 NBD1 (gray). A clear salt bridge is formed between D510 and K564. Panel B shows local change in NBD1 structure at V510 loop (in grey) with the addition of the V510D mutation (blue). WT NBD 1 crystal structure shown in green for positional reference. Panel C/D K564 moves about 5.4Å between the standard and V510D conformations. It appears to alternate from the standard rotamer to a new rotamer capable of creating a salt bridge with D510. In chain A (at left), K564 is mostly in the salt-bridge conformation and partially in the standard NBD1-1D conformation. In chain B (at right), K564 is fully in the salt-bridge conformation. Density looks better for the salt-bridge conformation, but there is a population in the standard conformation.

The observed interaction of the D510 side chain with K564 in the crystal structure supports the hypothesis from our MD simulations that D510 predominantly interacts with residues in NBD1 to exert its stabilizing effect on CFTR. Interaction with K564 may stabilize the F508 loop, which in turn would be expected to increase productive folding of NBD1 and proper assembly of the full–length channel, a hypothesis that is consistent with corresponding *in vitro data.* The mutation also eliminates the exposed hydrophobic side chain of V510, further improving NBD1 stability. Since R1070 is not present within NBD1, our isolated NBD1 structure cannot confirm potential interactions of D510 with R1070; however, given that we see a similar pattern of effects for the V510D and K564 mutants in the context of full-length CFTR and NBD1, it appears possible that the predominant effect of V510D is confined to NBD1.

### ΔF508–CFTR Helical Unraveling

An interesting observation from our ΔF508 CFTR MD simulations was the apparent instability of the TMD2 domain and a propensity of TM10 and TM11 to lose helical structure as a consequence of the ΔF508 mutation. These significant structural changes in helices 10 and 11 (Figure 3) may contribute to the short half-life and dysfunction of ΔF508-CFTR. Moreover, when we evaluated ΔF508/V510D-CFTR using the same parameters, the model suggests that the structural integrity of these domains is somewhat restored. The MD simulation results for ΔF508/R1070W-CFTR suggest that the resulting helical stabilization for this SSSM is less than that observed for ΔF508/V510D-CFTR.

In an attempt to validate the helical unraveling of ΔF508-CFTR predicted by the MD simulation and to better understand the impact ΔF508 might be having on TMD2, including whether NBD1 stabilization with and without ICL4 correction might lessen this impact, we created a series of constructs introducing mutations along TMs 10 and 11 with the potential to stabilize helical structure. Informed by our models, we selected residues Q1042, P1050 and L1096 (Figure 8), all within the region predicted to unravel, based on their likelihood to create salt bridges with existing residues in close proximity, and replaced each with charged residues in WT, ΔF508-CFTR, and ΔF508/V510D-CFTR expression constructs. To ensure that the potential introduction of salt bridges along TMs 10 and 11 did not interrupt normal CFTR trafficking, we first tested the mutations on the WT-CFTR background and saw no changes to normal CFTR maturation (data not shown). We then evaluated these mutations for their impact on ΔF508- and ΔF508/V510D-CFTR. No measurable increase in CFTR complex glycosylation and maturation was observed when the charged residue mutations were added to ΔF508-CFTR, apart from the P1050R mutation (data not shown). CFTR complex glycosylation was significantly augmented when V510D was added to ΔF508-NBD1; an increase in fully glycosylated CFTR was seen for several salt-bridge mutations when added to ΔF508/V510D-CFTR, in particular for mutants P1050K and P1050R (Figure 8). Based on these results, P1050R was identified as a potentially strong second-site suppressor of ΔF508-CFTR helical unraveling and was marked for further evaluation

**Figure 8.**
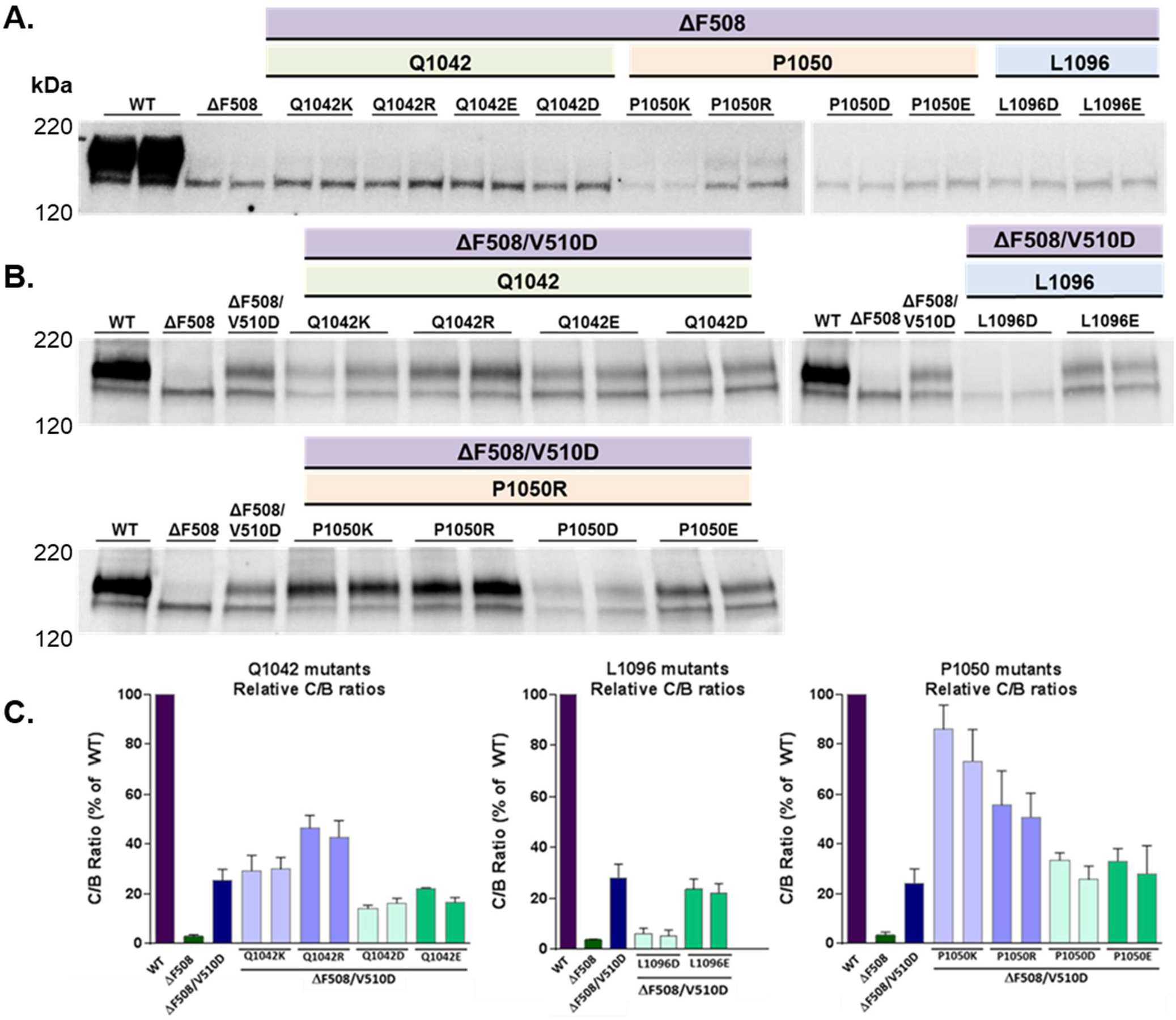
Western blot and C/B ratio analysis of helical stabilizing mutations along TMs 10 and 11. Initial (A) and repeat (B) analysis of ΔF508/V510D CFTR expression with the addition of charged residue mutations at P1050 along TMs 10 and 11 confirm the stabilizing effect of P1050R. Expression was normalized using a bicistronic GFP expression control. Maturation is increased in nearly all cases when stabilizing mutations are added to ΔF508/V510D CFTR as measured by an increase in fully glycosylated CFTR. P1050R was identified as a strong stabilizer of ΔF508/V510D CFTR. A representative western blot is shown comparing each of the four mutations evaluated, with samples loaded onto the gel as biological replicates. Corresponding C/B ratio values (C) reflect an n of 3 separate experiments for each mutational replicate.

At Q1042 and P1050, the increase in C-band appeared greater when positively charged basic amino acids Lys or Arg residues were swapped in, compared to the negatively charged Asp and Glu residues, and the slightly larger Arg resulted in the greatest level of CFTR maturation in all cases. Additionally, when such mutations were paired with V510D, a synergistic effect was seen on the levels of CFTR maturation when the double mutant was compared to either ΔF508-CFTR or ΔF508/V510D-CFTR alone. We observed that when the P1050R mutation was made to ΔF508/V510D-CFTR it restored trafficking better than V510D alone, and in a few instances, restored trafficking to near WT levels (Figure 8). This mutation was subsequently used for more in-depth analysis of helical stabilization. While P1050K had the highest C/B ratio by densitometry analysis, it produced slightly less CFTR overall.

#### The Role of V510D in P1050R-Mediated Helical Rescue

The V510D suppressor mutation is postulated to correct ΔF508-CFTR dysfunction through partially independent effects on the stability of both NBD1 and the ICL4-NBD1 interface (20). We sought to understand the extent to which the apparent effects of V510D (see above) on helix stabilization were dependent on each mechanism. To accomplish this, the V510D interaction with R1070 at the ICL4 interface was disrupted by adding the R1070S mutation to ΔF508/V510D-CFTR, both alone and in combination with the TM10 salt-bridge mutation P1050R.

As mentioned above, when P1050R was added to ΔF508-CFTR, a small improvement in CFTR maturation occurred that was approximately half that of the ΔF508/V510D mutation alone (Figure 9). A synergistic increase in trafficking and maturation was measured when P1050R was combined with V510D, which restored the level of mature CFTR at the cell surface to ~80% of WT levels when analyzed by both western blotting (C to B ratio) and HRP trafficking assays (Figure 9). When the ICL4 interface was interrupted by the addition of R1070S, however, V510D-mediated rescue of ΔF508-CFTR decreased from ~15% to 5% of WT levels. A similarly modest rescue effect (~5% of WT) was seen when P1050R was added to ΔF508/R1070S-CFTR; however, a synergistic effect was seen when both P1050R helical stabilization and V510D NBD1/ICL4 stabilization were added to ΔF508/R1070S-CFTR, which improved trafficking to roughly half that of WT CFTR. To evaluate NBD1:ICL4 interface stabilization with and without P1050R, we assessed the impact of R1070W on ΔF508-CFTR and saw a very modest improvement in trafficking similar to V510D alone, as anticipated. When ΔF508/R1070W/P1050R was evaluated, trafficking was restored to about half that of WT-CFTR, similar to the combination of V510D and R1070S with P1050R.

**Figure 9.**
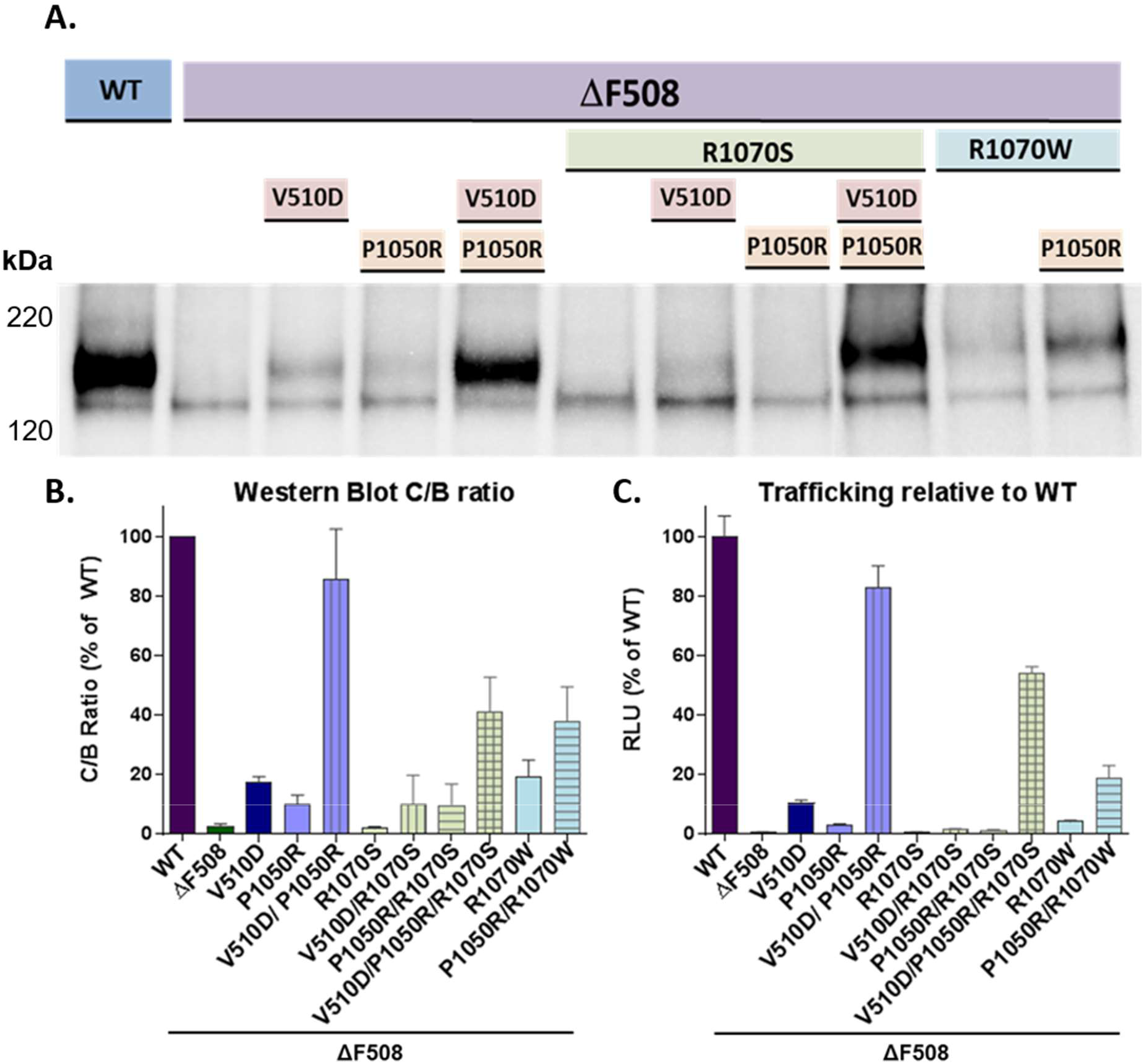
The impact of ICL4 and NBD1 stabilization on P1050R-mediated rescue of ΔF508-CFTR was evaluated by western blotting (A), the corresponding C/B ratio graph (B) and HRP trafficking (C). Adding P1050R to ΔF508-CFTR results in very modest improvements in CFTR maturation. However, when suppressor mutation V510D is paired with P1050R, the greatest impact on ΔF508-CFTR maturation and trafficking is seen at about 80% of WT CFTR. When R1070S is then added to explore the impact of interrupting the NBD1:ICL4 interface, this level of CFTR maturation drops from ~80% to 50%. When R1070W is included to partially restore the ICL4 interface and V510D is removed, the impact on ΔF508-CFTR is about half that of V510D, and the addition of P1050R to R1070W/ΔF508-CFTR only improves this to roughly 25%, suggesting that NBD1 stabilization is critical to the P1050R rescue effect of ΔF508-CFTR. For western blot, n=3; for trafficking assay, n=6 for each assay plate, with 3 replicate assays performed.

Next, we disrupted V510D-mediated NBD1 stability by again introducing K564A to remove the V510D salt-bridge interaction. Consistent with earlier results, V510D rescue of ΔF508-CFTR was diminished by the replacement of a charged lysine residue at 564 with an alanine. However, when P1050R was added to ΔF508/V510D/K564A-CFTR, a synergistic effect was again seen and trafficking was restored to ~40% of WT levels, despite the loss of K564 (Figure 10), supporting the hypothesis that V510D requires K564 for complete NBD1 stabilization, which is an essential component of ΔF508-CFTR maturation and trafficking.

**Figure 10.**
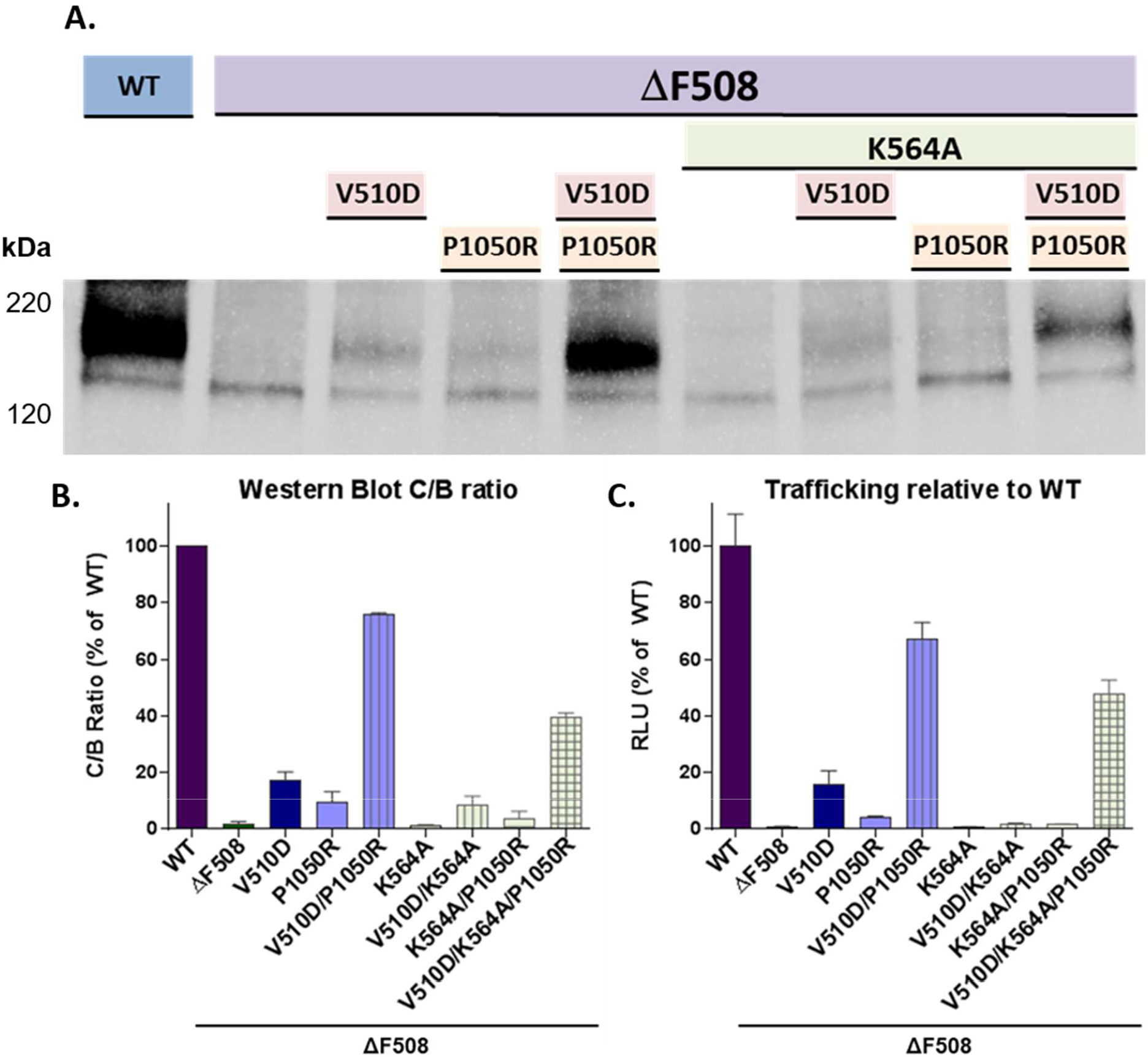
NBD1 stabilization and K564/V510D interactions are critical for P1050R rescue of ΔF508. K564A leads to a 30-40% reduction in V510D/P1050R rescue of ΔF508-CFTR as measured by immunoblotting (A & B) and HRP trafficking assays (C), underscoring the importance of NBD1 stabilization on P1050R ΔF508-CFTR rescue. For western blot, n=4; for trafficking assay, n=8 for each assay, with 3 replicate assays performed.

To understand whether the element of P1050R’s mechanism that is NBD1-dependent required V510D specifically, we replaced V510D with alternate NBD1 stabilizing mutations F494N/Q637R (2S), or F494N/Q637R/F429S (3S) (1,5,34), which do not reside at the ICL4 interface. When the 2S or 3S mutations were added to ΔF508-CFTR, a modest amount of rescue occurred, as evidenced by western blotting and HRP trafficking (6.5% and 9.5% of WT, respectively, Figure 11). However, when the NBD1 stabilizing 2S or 3S mutations were coupled with P1050R, maturation and trafficking of the resulting protein improved synergistically, with ΔF508/P1050R/2S reaching ~40% of WT and ΔF508/P1050R/3S resulting in ~80% of WT trafficking levels

**Figure 11.**
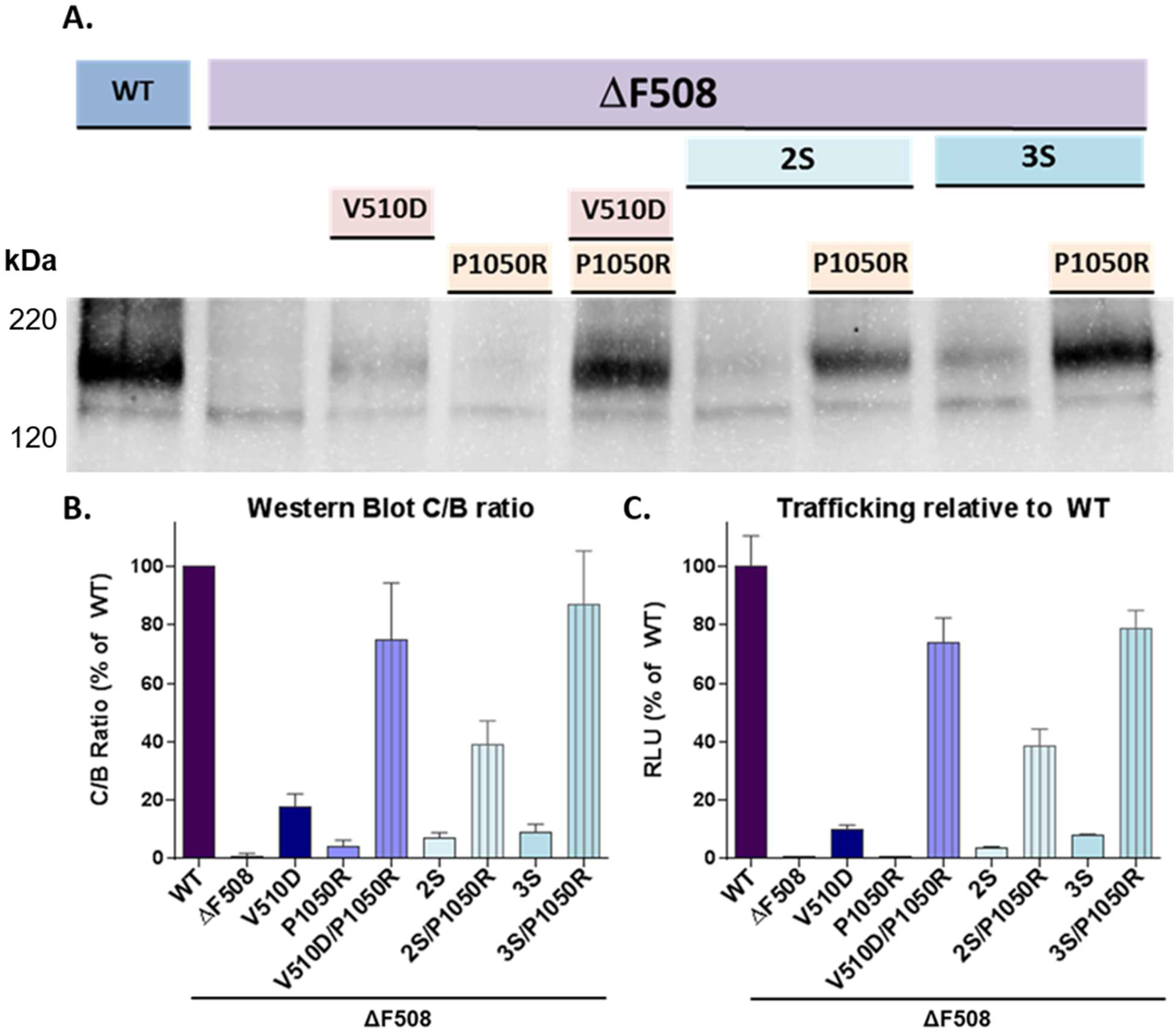
The role of NBD1 stabilization on ΔF508-CFTR rescue does not require dual correction by V510D when P1050R helical stabilization is added. When the 2S or 3S mutations were added to ΔF508-CFTR, improvements in trafficking of 6.5% and 9.5% of WT respectively were seen. However, when 2S or 3S mutations were coupled with ICL4 stabilizing mutation P1050R, maturation and trafficking significantly improved. ΔF508/P1050R/2S resulted in ~40% of WT and ΔF508/P1050R/3S in ~80% of WT trafficking levels. Western blot analysis (A) and C/B ratios (B) represent an n=4; for trafficking assay (C), n=8 for each assay, with 3 replicate assays performed.

### Functional Impact of the V510D and P1050R Stabilizing Mutations on ΔF508-CFTR

Electrophysiology was performed on a subset of the CFTR variants to understand more thoroughly the effect of the V510D (+/− K564A) and P1050R mutations on chloride channel function as compared to WT and ΔF508-CFTR (Figure 12). For the initial set of functional analyses, Fischer rat thyroid (FRT) cells were transiently transfected with DNA expressing WT-CFTR, ΔF508–CFTR, and ΔF508/V510D–CFTR with and without K564A. The maximal current after the addition of both Forskolin and CFTR potentiator Genistein (FP_max_) for each tested variant was then compared to WT and ΔF508-CFTR. Data trends were similar to those of the HRP trafficking and western blot assays (Figure 5).

**Figure 12.**
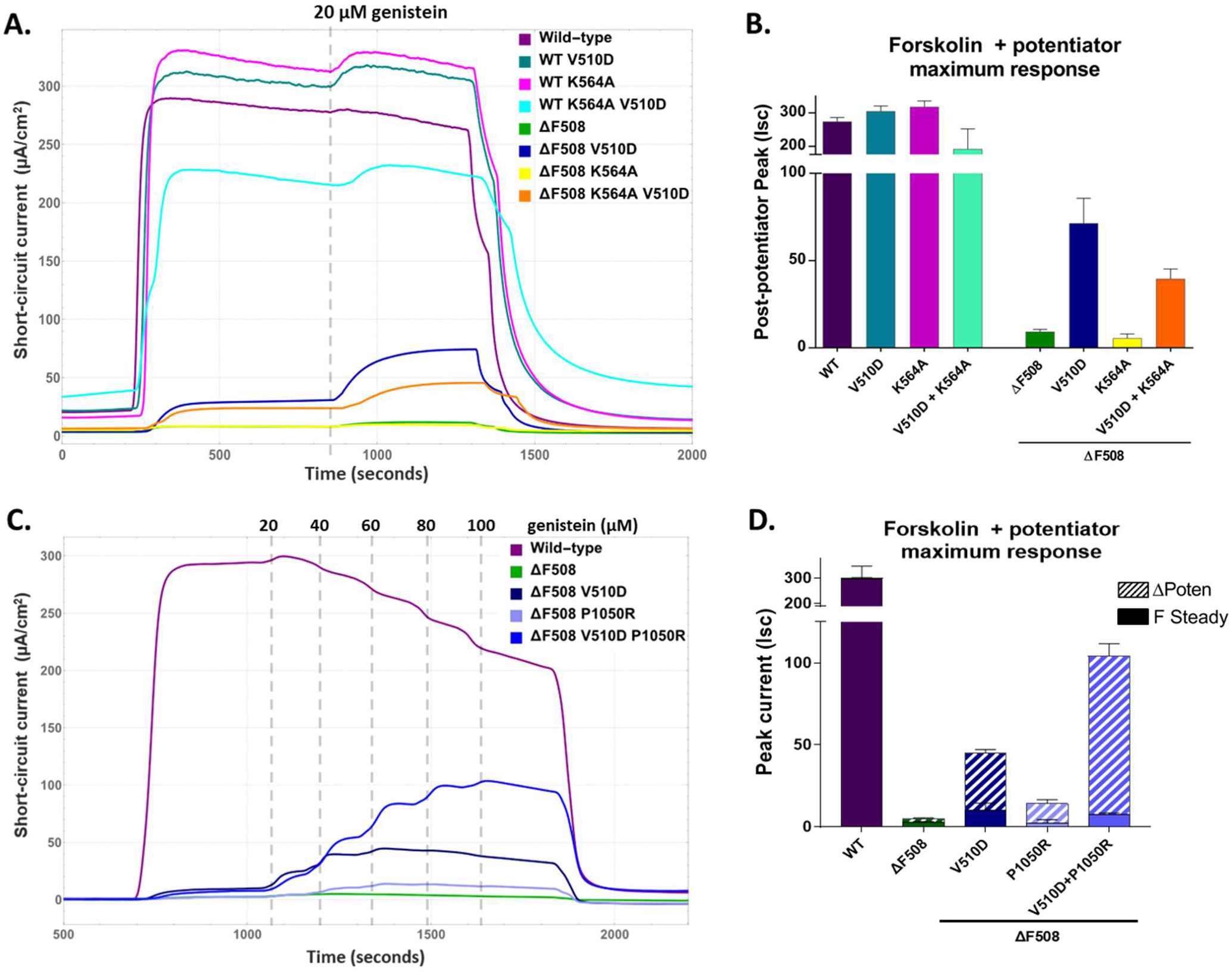
Electrophysiological properties of ΔF508/V510D CFTR +/− K564A or P1050R. Fischer rat thyroid cells expressing mutated CFTR constructs were evaluated for channel conductance to determine how V510D impacts ΔF508-CFTR function in the presence and absence of proximal residue K564A (A & B) or ICL4 stabilizer P1050R (C & D). In all cases, cells were treated with sequential application of 10 μM total forskolin (Fsk), 20 μM genistein as single (A & B) or repeat (C & D) dosing, and 20 μM CFTR-inh 172. Panels include (A) current traces for V510D/K564A mutants on a WT or ΔF508-CFTR background; (B) quantitative comparison of constructs displayed as peak forskolin + potentiator max response; (C) current traces for ΔF508/V510D +/− P1050R compared to WT-CFTR, and (D) comparison of total max chloride conductance in response to Fsk (denoted as F Steady) plus additive current from peak potentiation with 100 μM genistein (denoted as ΔPoten).

Additional analyses were then performed on P1050R-containing CFTR mutants with and without V510D. The initial maximal current for ΔF508/V510D/P1050R-CFTR post-Forskolin addition was similar to ΔF508/V510D-CFTR alone, suggesting that despite meaningful increases in CFTR trafficking and maturation, P1050R may hinder normal channel function. To determine whether this impaired CFTR could be rescued via potentiation, genistein was added to all chambers in 20 μM increments to a final concentration of 100 μM. Genistein addition resulted in a 12-fold dose-dependent increase in current for cells expressing ΔF508/V510D/P1050R-CFTR, bringing the average FPmax from ~7.7 Isc to 96.4. Interestingly, this level of genistein addition also led to significant increases in channel conductance for cells expressing ΔF508/V510D-CFTR and ΔF508/P1050R-CFTR and dose-dependently interfered with normal function in WT at higher concentrations (Figure 12).

## Discussion

This work was intended to accomplish three goals: i) to gain insight into the effects of the ΔF508 mutation on CFTR conformational stability, ii) to determine how second-site suppressor mutations may influence these, and iii) to validate a molecular model of ΔF508-CFTR (lacking R-domain), all in the absence of a full-length structure.

Given the potential impact of structure-based drug design on the development of CF therapeutics, the structural and biophysical properties of full-length CFTR continue to be of great importance. Recently, multiple high-resolution cryo-EM structures of CFTR were published, providing “snapshots” of both inward- and outward-facing conformations, as well as new insights into CFTR channel gating and function (24,25,27). Ultimately, however, the native structure of ΔF508-CFTR and a better understanding of the mutation’s impact on CFTR conformational dynamics would be invaluable yet remains challenging given the protein’s instability. And while a high-resolution structure of ΔF508-CFTR with stabilizing mutations might provide important information, this stabilization may also mask key elements of the ΔF508 folding defect. *In silico* models and MD simulations may offer a complementary view of the atomic-level interactions within the protein, as well as some understanding of the ΔF508 mutation’s impact on CFTR structure and how second-site suppressor mutations may influence this. In addition, it helps us develop testable hypotheses that, ultimately, could guide the discovery new therapeutics.

One aspect of the CFTR and Sav1866 structures that is notably different is the diameter of the channel lumen, an observation highlighted by Zhang *et al.* (23) when comparing the structure of the homologous protein Sav1866 to phosphorylated zebrafish CFTR (PDB ID: 5W81). In Sav1866, the diameter of the channel lumen that results from the separation of TM 4/6 and TM 10/12 is relatively large so as to accommodate drug-like substrates (23). In contrast, the diameter of the CFTR channel lumen is considerably smaller but is of sufficient size to provide a conduit for chloride ions. Because of this difference, several TM helices, which in our CFTR model were based on Sav1866, are rotated and displaced relative to the recently published CFTR cryo-EM structure (27). Notably, such differences are mostly limited to the extracellular side of the CFTR intramembrane region. Perhaps surprisingly, Sav1866 served as an appropriate template for the CFTR intracellular TMs, where substantial structural similarity is seen between the model and the solved structure (Figure 1). CFTR’s NBD1 and NBD2 (Figure 1; I and J), were modeled from crystal structures of the isolated domains. Both are very similar to the CFTR cryo-EM structure, including side chain orientations and, importantly, their interactions with ICL regions. This similarity may not be surprising, as the cryo-EM structures and our homology model incorporated the same NBD1 and NBD2 crystal structures as starting models for these domains.

While the results from our MD simulations are for a single random seed, and multiple runs with different random seeds would be required for a comprehensive statistical analysis, the data gathered provided a good starting point for hypothesis generation and subsequent experimental verification. The secondary structure analysis that resulted was in close agreement with previously reported experimental data for WT- and ΔF508-CFTR stability with and without V510D or R1070W (20,29,35,36). Our simulation also provided a solid platform for testable hypotheses regarding the impact of ΔF508 on TMD2 and critical interactions that support V510D suppression of ΔF508, which we then successfully verified experimentally. For example, our models clearly show stronger conservation of ΔF508-CFTR secondary structure with the addition of V510D compared to R1070W, an outcome previously predicted to be the result of V510D’s dual stabilization of both the NBD1:ICL4 interface and NBD1 alone, as opposed to R1070W’s primary role of interface correction (6,37).

The impact of these mutations alone and with additional NBD1 stabilizers in full-length CFTR, coupled with structural and stability data collected from solubilized ΔF508/V510D-NBD1, support the hypothesis that V510D plays a role in both interface correction and NBD1 stabilization, an outcome we observed in both WT and ΔF508-CFTR. Given that no reduction in CFTR maturation, trafficking or function was seen when residue K564 was altered within the context of WT-CFTR, regardless of the presence of V510D, it is likely that in WT/V510D-CFTR, no interaction exists between D510 and K564 (Figures 4 and 5). Rather, D510’s primary interaction becomes the salt-bridge with R1070, creating an interface that is too rigid, and may hinder the flexibility seen when the interface is mediated by F508. This level of interaction is not seen in ΔF508-CFTR, however, likely due to the flexibility and solvent exposure of the V510 loop, which may make this interaction easier to disrupt.

### Presence and Position of Residue K564 is Key to V510D Suppression of the ΔF508 Trafficking Defect

In addition to providing support for V510D’s dual role of ICL4 and NBD1 stabilization, we experimentally validated the salt-bridge interaction between K564 and V510D that was predicted from our MD simulation of ΔF508/V510D-CFTR in several different ways. Such support includes quantitative data collected from CFTR immunoblotting, HRP-trafficking and electrophysiology experiments, which were highly correlative (Figures 4 and 5). In all cases, the addition of V510D resulted in modest rescue of CFTR trafficking (an average of 5-10% of WT), which was diminished by half with the removal of K564’s positive side chain. The near-total loss of trafficking that occurred in the case of K564S, (regardless of V510D; Figure 5) may be attributed to perturbations arising from the addition of a serine hydroxyl group to the tail-end of the α-helix spanning residues 549-564 within NBD 1. In such a conformation, it is energetically unfavorable for the hydroxyl group to lack a hydrogen bond interaction and, thus, may compete with the protein backbone for available hydrogen bonds, potentially destabilizing the domain thus altering its conformation.

Another possible explanation for the inactivity of K564S is the potential creation of a phosphorylation site for Casein Kinase 2. The Ser phosphorylation motif for CK2 kinase is a critical acidic residue (preferably Asp) at position +3 (38–40). Phosphorylation potential is enhanced further if it has additional acidic residues (again, preferably Asp) spanning from −2 to +7 (38,41–43). Additionally, this requires the absence of a proline or bulky hydrophobic side chain at n+1, or bulky hydrophobic doublets at +1 and +2 (43,44). All of these criteria are met in CFTR sequence 564 – 572, i.e. **K564(S)-D-A-D-L-Y-L-L-D.** Ser564 is at the C-terminal end of an α-helix while the N-terminal end and the preceding loop have close interactions with ATP; a number of stabilizing residues are located in this region and hence any perturbation of this region may destabilize NBD1 and may have functional consequences due to its interaction with ATP.

Despite the modest occurrence of an interaction that appeared between V510D and R487 in our salt bridge analysis performed for the ΔF508/V510D-CFTR MD simulation, mutating the positively charged arginine residue at position 487 to an alanine does appear to significantly reduce trafficking and function of both WT- and ΔF508-CFTR (data not shown). This may be due to the residue’s potential role in ATP binding and hydrolysis, specifically transition state stabilization [44]. Indeed, no interaction is observed between D510 and R487 in our crystal structure. However, we see that R487 is involved in a crystal contact in our crystal form, which may make it unavailable for interaction with D510 in that simplified system. As a result, a high-resolution structure of full-length ΔF508/V510D CFTR may be required to determine whether this predicted interaction with R487 might contribute to V510D’s activity as a SSSM.

### The Occurrence and Subsequent Rescue of TMD2 Helical Unraveling in ΔF508-CFTR

Our MD simulations data also suggested that TMD2 helical unraveling may be a key component of ΔF508-CFTR destabilization, a prediction that has not previously been reported. Our models suggest that the α-helical region of ICL4 spanning residues 1034 to 1050 (aa sequence: ESEGRSP) is one of the most destabilized within ΔF508-CFTR, likely owing to the presence of helix-breaking residues serine and proline, which make this region vulnerable to conformational perturbations. In WT-CFTR, when the NBD1:ICL4 interface and the α-helical conformation of this region are intact, the model predicts that salt bridges exist between E1044 and K1041, and potentially with R1158 or K978 (from ICL2), which may help stabilize the loop. In ΔF508-CFTR the model predicts that these salt bridges are broken, as E1044 and K1041 move away from each other, and L1040, which may normally create stabilizing interactions with L1091 and W1089 in TM11, becomes completely exposed. Based on this information, we hypothesized that the addition of polar residues at the intracellular side of helices 10 and 11 might produce stabilizing salt bridge interactions, effectively creating second-site suppressors of ΔF508.

To determine which mutation sites would produce the most impactful stabilization, we referred to sequence and structural information for TMD1 for guidance, given its presumably greater stability that our model suggests. Sequence comparison of the two TMDs (Figure S6) shows a higher prevalence of charged residues within TMD1 (aa 66-431), which contains 19 Arg and 22 Lys, as well as 21 Glu and 9 Asp for a total of 41 positively charged and 30 negatively charged side chains. By comparison, TMD2 (aa 845-1198) contains only 28 positively charged side chains (13 Arg and 15 Lys), and 18 negatively charged (9 Asp and 9 Glu). (We did not take into account His residues in either case, as they are normally neutral.) Based on this comparison, three residues were highlighted by our analysis as potential stabilizing mutation sites: Q1042 and P1050, which are both on TM10 of ICL4, and L1096, which resides at the bilayer interface of TM11, with preference given to the former two based on their likelihood for success. In all cases, polar residue insertions made no improvement to CFTR maturation and trafficking when added to ΔF508-CFTR (Figure 8), which is unsurprising given the unstable nature of the protein overall, and if our model is any indication, the significant helical disorder that may be present. However, when the charged residue substitutions are paired with V510D, we see stabilizing trends emerge. Overall, a greater increase in CFTR trafficking was seen when replacements were made at Q1042 and P1050 with basic residues Arg and Lys, suggesting the possible interaction of either residue with E1046; however, a favorable position of the sidechains would be required. Moreover, when compared to Lys, Arg substitutions appeared to result in slightly higher levels of C-band, perhaps owing to the presence of a guanidinium group, which may enable a greater number of CFTR-stabilizing interactions (45). The replacement of either Q1042 or P1050 with acidic residues Asp and Glu did not appear to improve trafficking more so than V510D alone, suggesting that the insertion of a negative charge, while not potentially destabilizing, provided no suppression of helical instability. This is in keeping with the “positive inside rule,” which states that positively charged residues (Arg and Lys) will frequently be located at the cytoplasmic edge of transmembrane helices, (46,47), with the opposite being true for negatively charged residues (48), a trend that is supported by extensive statistical observations for most membrane proteins (48,49). The proposed utility of this “charged-residue flanking bias” (48) is that positively charged amino acids bordering hydrophobic helices might be involved in TM orientation within the lipid bilayer (47), as well as helix-helix interactions (47,50,51). With regard to L1096, we must consider that leucine is highly hydrophobic, and is known to play a critical role in both helical stabilization and TM helix-helix interactions (47,48,52). Unsurprisingly, modifying this residue did not improve ΔF508-CFTR trafficking as shown by western blot (Figure 8), and in the case of the L1096D mutation, abolished the V510D stabilizing effect.

### Synergy Between NBD1 Stabilizing SSSMs and P1050R

The impact of V510D on NBD1 and interface stabilization contributes both to its activity as an SSSM and its synergy with P1050R. Our studies with NBD1 stabilizing mutations help us understand the relative importance of these two effects. As described above (Figure 9), NBD1 stabilizing mutations F494N/Q637R (2S), or F494N/Q637R/F429S (3S) (1,5,34) both robustly synergize with P1050R to further rescue ΔF508-CFTR maturation and trafficking in the absence of V510D. The 2S mutations, which together stabilize purified ΔF508-NBD1 by ~2 °C (53), rescue ΔF508-CFTR maturation to ~40% of WT levels when combined with P1050R. A larger impact is seen in the presence of the 3S mutations, which together stabilize purified ΔF508-NBD1 by 5.7°C (5). Rescue of the ΔF508-CFTR maturation and trafficking are highly similar when either 3S/P1050R or V510D/P1050R is present. In each case, rescue to levels >80% of WT was observed. It is noteworthy that the 2.5 to 3°C increase in ΔF508-NBD1 stability when V510D is present is far closer to that seen with 2S than with 3S. The ΔF508-CFTR rescue by V510D in the presence of P1050R is therefore greater than would be predicted from its impact on NBD1 stability alone. It is also noteworthy that R1070W/P1050R rescues ΔF508-CFTR maturation to a similar level, viz 40% of WT, seen in the presence of 2S/P1050R. These levels of ΔF508-CFTR rescue are roughly half those provided by P1050R in the presence of V510D, consistent with its dual effect. These findings are also consistent with our studies of ΔF508/R1070S/V510D CFTR, where, relative to ΔF508/V510D CFTR, a significant reduction in apically localized CFTR and the band C/band B ratio are observed. Similar differences in relative rescue levels are seen for these mutations in the presence of P1050R (Figure 9).

Despite the ability of P1050R to increase ΔF508 CFTR maturation and trafficking in the presence of V510D, the rescued channel appears to possess impaired function. As shown in Figure 12C, expression of CFTR channel function in rescued ΔF508/V510D/P1050R increases significantly upon treatment with the CFTR potentiator genistein, even at higher genistein concentrations, which have been shown to reduce WT CFTR function (54,55). Such findings may have implications for therapeutic CFTR correction, as it suggests pharmacologic agents that might increase TM10 stability, could also yield ΔF508-CFTR that is functionally deficient, and that combination with a CFTR potentiator could be needed to rescue activity.

## Summary

Together, our findings further highlight the dual nature of the V510D stabilizing mechanism. Our experimental data validate novel predictions derived from our *in silico* CFTR model and MD simulations of it. The data provide evidence that V510D and K564 have additive effects on ΔF508-CFTR rescue, which also supports the requirement of K564 for V510D stabilization, as seen from the MD simulations and subsequent structural biology studies. Central among our validated predictions are: (i) the previously undescribed impact of ΔF508 on the stability of TMD2 helices, (ii) the contribution of helical unravelling to ΔF508-CFTR dysfunction, and (iii) the existence of novel SSSMs within TM10 that partially counteract this aspect of the ΔF508 defect. The data demonstrate that the novel SSSM P1050R acts by a mechanism distinct from those of R1070W, V510D or NBD1 stabilizing mutations. Indeed, P1050R’s synergy with interface and NBD1 stabilizing SSSMs underscores the potential therapeutic value of small molecule ΔF508-CFTR correctors that share its presumed mechanism of action, namely TM10 stabilization.

## Materials and Methods

### Homology Modeling

Homology models of full-length WT, ΔF508, ΔF508/V510D and ΔF508/R1070W CFTR were generated with Discovery Studio Software package (56) using the crystal structure of SAV1866 in the outward facing conformation (PDB ID: 2HYD) as a template. We replaced the SAV1866 NBD1 domain with the CFTR NBD1 crystal structure (PDB ID: 2PZE) and made the appropriate corrections for all missing side chains and residue substitutions. Likewise, the ΔF508-NBD1 crystal structure (PDB ID: 2PZF) was incorporated for the ΔF508-CFTR variant models. For modeling NBD2, the crystal structure of NBD2 fused to maltose-binding protein (PDB ID: 3GD7) was used as the template. Relative positioning of NBD1 and NBD2 were based on Sav1866 and NBD1 homodimer structures. The R-domain, which ranges from approximately residues 641-849, was not included in our model, as no homologous template was available for this domain. To create the ΔF508 SSSM variants, stabilizing mutations V510D and R1070W were individually added to the ΔF508-CFTR model, and any resulting steric hindrances were relieved by local energy minimization. All models were further refined using Maestro Protein Preparation Wizard (57).

### Molecular Dynamics Simulations

We used the Desmond System Builder (58) to place a pre-defined lipid bilayer of 1-palmitoyl-2-oleoyl-sn-glycero-3-phosphocholine (POPC) using transmembrane boundaries reported in Uniprot (59) for all homology models. Each system was solvated with a TIP3P (transferable intermolecular potential with 3 points) water model (Jorgensen 1983) and neutralized with chlorine counter ions. We also added sodium chloride at a concentration of 0.15 M, leading to roughly 160K atom systems. The system was then equilibrated using the defaults settings of the Desmond package. After equilibration, all simulations were carried out using the Desmond-GPU package implemented in Maestro from Schrodinger (60) at a constant-temperature, constant-pressure ensemble (NPT) of 300 K and 1.01 Bar, respectively, for a time of 1 μs and a frame-capture interval of 800 ps resulting in a total of 1,251 frames. Subsequent analyses were performed using Maestro (60), VMD (Visual Molecular Dynamics) (61), MOE™ (Molecular Operating Environment) from Chemical Computing Group (62) and several modules that were developed in-house using Python, SVL and Perl scripting languages.

For each model, event analysis was completed to identify changes in secondary structure, residue surface area/volume/geometry, variation in residency time of chosen residues in select regions of the protein, and the presence or absence of salt bridges (63). Residues of interest for secondary structure analysis included neighboring regions S495-E514 within NBD1 and Q1035-R1102 at ICL4. Solvent accessibility calculations were performed using the Euclidean Distance Transform surface calculation method (64)

### Construct Generation and Expression of Full-Length CFTR Mutants

#### CFTR Plasmid Mutagenesis

QuikChange Single and Multi-Site Directed Mutagenesis system (Agilent Technologies) was used to modify select residues located in NBD1 and along helices 10 and 11 in TMD2. Within NBD1, K564A/S and R487A/S mutations were made, and along TM10 and TM11, charged residues were introduced at Q1042, P1050 and L1096 in full-length WT, ΔF508–CFTR, and ΔF508/V510D-CFTR expression constructs. A second round of constructs was made to include suppressor mutations F494N, Q637R and F429S or R1070W alongside P1050R +/− V510D in ΔF508-CFTR. For some mutants, a duplicate set of mutated WT and ΔF508-CFTR plasmids that express an HRP tag on ECL4 (between S902 and Y903) was created to evaluate trafficking (see Figure 2.9D.). All constructs contained a “self-cleaving” GFP transfection control, introduced by 2A-based bicistronic expression. In all cases, mutations were verified by Sanger sequencing.

#### CFSMEo-Cell Culture

CF patient-derived CFSME_o-_ cells (kindly provided by Dr. Dieter C. Gruenert, UCSF) were maintained at 37°C in Minimum Essential Medium with Earle’s salts, supplemented with 10% (v/v) FBS, 2 mM L-glutamine and 1% (v/v) pen/strep on ECM-coated flasks (10 μg/mL human fibronectin, 30 μg/mL bovine collagen I, 0.1% BSA in LHC basal medium). For evaluation of CFTR trafficking, cells were plated on collagen-coated plates (Corning BioCoat multiwall plates, Corning, Inc) and transiently transfected using FuGENE HD transfection reagent (Promega US, Madison, WI) according to the manufacturer’s instructions.

#### Western Blotting

CFSME_o-_ cells were plated at 200,000 cells per well in a 6-well, collagen-coated plate and transfected with 3 μg CFTR variant plasmid DNA per well. After 48 hours, cells were washed twice with PBS and collected in cold lysis buffer (25 mM Tris-HCl pH 7.4, 150 mM NaCl, 1 mM EDTA, 1% NP-40 and 5% glycerol) supplemented with protease inhibitors (Roche Complete EDTA-free). Protein concentrations were determined by bicinchoninic acid assay (BCA protein assay kit, Pierce Thermo Fisher Scientific). Whole cell lysates were separated on 7.5% Criterion TGX 7.5% SDS-PAGE gels (Bio-Rad) and transferred to nitrocellulose membranes using a semi-dry transfer apparatus at 20 V for 10 minutes. Membranes were probed with anti-CFTR monoclonal antibody 570 (obtained from UNC Chapel Hill) at a 1:1000 dilution to determine CFTR protein expression and maturation of each mutant construct. Differences in trafficking were measured by comparing the ratio of “C” band to “B” band by densitometry.

#### HRP-Trafficking

CFSME_o-_ cells were plated at 8,000 cells per well in 96-well collagen-coated plates and transiently transfected with CFTR variant plasmids containing an HRP reporter as described above. After 48 hours, cell culture media was removed, cells were washed with PBS, Luminata Forte HRP substrate (EMD Millipore, Burlington, MA) was added to each well, and the HRP reporter chemiluminescent signal on ECL4 was measured using an EnVision plate reader. When CFTR variants traffic to the apical cell surface, the HRP tag is exposed on the outside of the plasma membrane, rendering it accessible to HRP substrate. To ensure the measured signal was the result of extracellular HRP localization, a qualification experiment was performed using Brefeldin A, a lactone antiviral that inhibits protein transport from the endoplasmic reticulum to the Golgi apparatus [28]. Addition of this compound to the cultured CFSME_o-_ cells interrupted normal trafficking of the CFTR HRP reporter to the outer membrane, which translated into a reduction in extracellular HRP signal that was comparable to that of ΔF508-CFTR. Relative CFTR surface localization could then be determined in live cells by measuring the resultant chemiluminescent signal [29, 30].

To ensure that samples were being properly compared for expressed CFTR in the HRP-trafficking and immunoblotting experiments, samples were normalized by eGFP expression based on RFU values of each sample lysate. For this, 20 μL of lysate was added in triplicate to a 384-well white-walled plate (ProxiPlate-384 Plus, shallow-well microplate, PerkinElmer, Waltham, MA) and read on an EnVision plate reader (PerkinElmer) using excitation/emission wavelengths of 480/535 nm, and an anti-GFP antibody was used during immunoblotting. For the HRP trafficking assay, a semi-quantitative value (RLU) corresponding to the level of CFTR maturation was normalized to GFP expression.

### NBD1 Analysis

#### Protein Production

Human CFTR NBD1 variants (residues 387–678 less 405–436 for ΔRI or 405-436/647-678 for ΔRI/ΔRE, with and without selected mutations) were expressed in BL21 *E. coli* cells as N-terminal His_10_-Smt3 fusion proteins (65) using a pET24a-derived expression vector. Cultures were grown, harvested and processed as previously described (34), and the protein was purified from the cell lysate using a nickel ion affinity column and a HiLoad 26/600 Superdex 200 column. The His_10_-Smt3 affinity tag was removed and the sample was again passed through a nickel ion affinity column and gel filtration column for further purification. The resulting protein was concentrated and stored at −80°C in buffer containing 50 mM Tris, 150 mM NaCl, 5 mM MgCl2, 12.5% w/v glycerol, 2 mM DTT, 2 mM ATP, pH 7.6. Final purity and concentration were determined by SDS-PAGE.

#### Differential Static Light Scattering (DSLS)

Purified recombinant NBD1 was produced at Sanofi using previously described methods (2,66). Thermal stability of various NBD1 isoforms was evaluated by DSLS using the Harbinger Stargazer-384 instrument (Epiphyte Three, Toronto, Canada). NBD1 protein was diluted to 0.2 mg/ml in S200 buffer (50 mM Tris-HCl, 150 mM NaCl, 5 mM MgCl2, 2 mM ATP, 2 mM DTT, pH 7.6) containing 1% glycerol, and 10 μL of protein solution was aliquoted into wells of a 384-well low-volume optical plate (Corning Inc., Corning, New York), and 10 μL of mineral oil was then overlaid onto the protein solution. Once in the Stargazer instrument, the plate was heated from 25°C to 70°C at a rate of 1 °C/min to facilitate protein unfolding and aggregation. Throughout the experiment, visible light was shone on the protein from below, and images of the light diffraction pattern for each well was captured from above the plate every 30 seconds. A linear regression curve was generated for each well, representing the increase in light scattering over time. By integrating the pixel intensity curve of each well, and plotting the total scattered light value against temperature, an inflection point representing the temperature of aggregation (T_agg_) was calculated for each sample. Final data curves were generated using Wolfram Mathematica by plotting the full raw data set

#### Differential Scanning Fluorimetry (DSF)

Nano differential scanning fluorimetry (nanoDSF) was performed to evaluate NBD1 thermal stability by measuring intrinsic tryptophan (and to a lesser extent, tyrosine) fluorescence as a measure of protein denaturation. Experiments were performed using the Prometheus NT.48 nanoDSF (NanoTemper Technologies, Germany). For this work, protein was diluted to 1 mg/mL in S200 buffer containing 1% glycerol. Approximately 10 μL of each diluted sample was loaded into nanoDSF grade high-sensitivity capillaries (NanoTemper) in triplicate and placed on the instrument’s capillary loading tray. The instrument temperature gradient was set to increase from 20°C to 95°C at a rate of 0.5°C/minute. Protein denaturation was detected at emission wavelengths of 330 and 350 nm, and a ratio of 350/330 fluorescence intensities was created for each time point collected. This ratio was then plotted against temperature, representing the transition from properly folded to fully denatured protein. The first derivative of the resulting sigmoidal curve was again plotted against temperature, with the extreme of the derivative curve displaying the melting temperature (Tm) for each sample.

#### Crystal Structure

##### Crystallization

CFTR NBD1 V510D was crystallized by mixing equal amounts of protein at 6 mg/mL with 0.1 M Tris pH 7.6, 28% (w/v) polyethylene glycol 10,000. Crystallization reagents were from Hampton Research, Aliso Viejo, CA. Crystallization was induced by streak-seeding using crystals grown in 0.1 M HEPES, pH 7.5, 25% (w/v) polyethylene glycol 550 monomethylether, and plates incubated at 4°C. Crystals grew over several days and were frozen by quick dip in 28% (w/v) PEG 10, 0.1 M Tris, supplemented with 25% (w/v) ethylene glycol.

##### Data collection, processing, structure determination and refinement

Data were collected at the Advanced Photon Source (APS Argonne, IL) and processed using HKL2000 (67). Molecular replacement was carried out using Phaser (68) of the CCP4 suite (69). The structures were refined using Phenix (70), followed by manual corrections in COOT (71). The structure was inspected and analyzed in COOT and PyMOL (72).

### Electrophysiology

Transfected Fischer rat thyroid (FRT) cells grown to confluence on 6-well “snapwell” transwell inserts (Corning) were mounted in Ussing Chambers (Physiologic Instruments). The apical solution contained 140 mM NaCl, 5 mM KCl, 2 mM CaCl_2_, 1 mM MgCl_2_, 10 mM HEPES, 10 mM glucose, pH 7.4. In the basolateral solution, NaCl was replaced with 140mM Na Gluconate. CFTR current was induced upon addition of 5μM forskolin to both the apical and basolateral chambers and augmented with the addition of the CFTR potentiator genistein. Channel recordings were terminated after adding 20uM CFTR inh-172. Transepithelial current (Isc), conductance (Gt), and voltage were measured using a multichannel voltage/current clamp VCCM8 system (Physiological Instruments) and recorded using the Acquire and Analyze 2.0 software (Physiologic Instruments). Final data curves were generated using Wolfram Mathematica by plotting the full raw data set.

### Data availability statement

The atomic coordinates and structure factors (**PDB ID: 6WBS**) for the crystal structure found in Figure 7 have been deposited in the Protein Data Bank, Research Collaboratory for Structural Bioinformatics, Rutgers University, New Brunswick, NJ (http://www.rcsb.org/). Release of the structure is pending publication of this manuscript

